# Precise manipulation of site and stoichiometry of capsid modification enables optimization of functional adeno-associated virus conjugates

**DOI:** 10.1101/2023.09.07.556719

**Authors:** Sarah B. Erickson, Quan Pham, Xiaofu Cao, Jake Glicksman, Rachel E. Kelemen, Seyed S. Shahraeini, Sebastian Bodkin, Zainab Kiyam, Abhishek Chatterjee

## Abstract

The ability to engineer adeno-associated virus (AAV) vectors for targeted infection of specific cell types is critically important to fully harness its potential of human gene therapy. A promising approach to achieve this objective involves chemically attaching retargeting ligands onto the virus capsid. Site-specific incorporation of a bioorthogonal noncanonical amino acid (ncAA) into the AAV capsid proteins provides a particularly attractive strategy to introduce such modifications with exquisite precision. In this study, we show that using ncAA mutagenesis, it is possible to systematically alter the attachment site of a retargeting ligand (cyclic-RGD) on the AAV capsid to create diverse conjugate architectures, and that the site of attachment heavily impacts the retargeting efficiency. We further demonstrate that the performance of these AAV conjugates is highly sensitive to the stoichiometry of capsid labeling (labels per capsid), with an intermediate labeling density (∼12 per capsid) providing optimal activity. Finally, we developed technology to precisely control the number of attachment sites per AAV capsid, by selectively incorporating a ncAA into the minor capsid proteins with high fidelity and efficiency, such that AAV-conjugates with varying stoichiometry can be synthesized in a homogeneous manner. Together, this platform provides unparalleled control over site and stoichiometry of capsid modification, which will enable the development of next-generation AAV vectors tailored with desirable attributes.

## Introduction

Over the last decade, AAV has emerged as the most promising vector for human gene therapy, enabling the development of clinically approved therapeutics such as Luxturna and Zolgensma.^*1-5*^ It offers numerous unique advantages that have contributed to its success, including a lack of pathogenicity, superior safety profile, mild immunogenicity, and long-term gene expression without genomic integration. However, the current difficulty with targeting AAV vectors to a desired cell/tissue type with good efficiency and selectivity is a major weakness of this strategy.^*6*^ Most AAV gene therapies currently under clinical trials rely on native serotypes, which show suboptimal tissue tropism/selectivity. Consequently, high clinical doses are typically needed to achieve therapeutic efficacy, which significantly increases the likelihood of adverse off-target effects, while contributing to exorbitant treatment costs that severely restrict their accessibility.

Several approaches have been explored to engineer AAV vectors for more targeted gene delivery.^*7-12*^ These approaches encompass the directed evolution of novel capsids with altered tropism;^*13-16*^ the strategic insertion of unique peptide sequences into permissive locations within capsid proteins,^*10*^ which either directly bind a new receptor, or can be enzymatically tagged with receptor targeting functionalities;^*17, 18*^ and the fusion of antibody fragments with the N-terminus of the minor capsid protein VP2.^*19-21*^ Although valuable, the directed evolution strategy is time and labor intensive, and frequently falls short in generating vectors with desired attributes. Due to the delicate nature of the AAV capsid, comprising a complex interwoven assembly of 60 multifunctional capsid proteins, methods like peptide insertion or antibody fusion are frequently susceptible to introducing unfavorable perturbations, negatively impacting viral assembly, entry, or trafficking. Not surprisingly, such modifications are tolerated at a very limited number of sites.

Chemical attachment of targeting ligands on the virus capsid has recently emerged as a promising alternative for engineering its cell type specificity. This can be achieved by targeting canonical amino acid side chains such as lysine or arginine on the virus capsid.^*22-24*^ However, this strategy intrinsically lacks site-specificity, and yields a heterogeneous mixture of AAV conjugates where the sites and stoichiometry of labeling vary widely. In contrast, the genetic code expansion (GCE) technology^*25-27*^ can be used to site-specifically incorporate a noncanonical amino acid (ncAA) with a bioorthogonal conjugation handle into proteins, which could be subsequently labeled with a high degree of selectivity.^*26, 28-31*^ We and others have previously shown the feasibility of site-specifically incorporating ncAAs into the AAV capsid, and their subsequent use to introduce precise chemical modifications.^*32-39*^ Using this strategy, it has been possible to attach retargeting ligands onto the capsid, which allows the virus to utilize a non-native receptor to gain entry into cells. For example, by attaching a cyclic-RGD (cRGD) ligand onto the AAV capsid using strain-promoted azide-alkyne click chemistry (SPAAC), we were able to retarget AAV2 to the αvβ3 integrin receptor, which is overexpressed in various cancer cells.^*32, 33*^

The ncAA mutagenesis technology offers an opportunity to create AAV conjugates with a remarkable degree of versatility. Certain ncAAs are tolerated well across many surface exposed sites on the AAV capsid, providing access to a vast array of potential attachment sites.^*32*^ However, how the site of attachment impacts the performance of AAV conjugates has not been systematically explored. Additionally, established ncAA mutagenesis strategies yield AAV particles with 60 ncAA residues, one in each of the 60 capsid proteins. The controlled modification of the resulting capsid can produce capsids with different stoichiometry of capsid labeling (labels per capsid). However, the relationship between the stoichiometry of labeling and the performance of the AAV conjugates remains unclear. In this report, we carefully explore how the attachment site and stoichiometry impact the performance of an AAV-cyclic-RGD (cRGD) conjugate. By attaching cRGD at each of the 10 consecutive positions in the AAV capsid proteins that comprise the spike region, we show that the attachment site strongly influences the retargeting efficiency of the resulting AAV conjugates. We also generated and tested a series of AAV-cRGD conjugates with an increasing stoichiometry of capsid labeling, which revealed that an intermediate degree of labeling per capsid (approximately 12 cRGDs per capsid) provides optimal retargeting efficiency. Furthermore, to enable the synthesis of such optimal low-stoichiometry conjugates in a homogeneous manner, we developed technology to introduce a ncAA into only the minor capsid proteins (either VP1 or VP2 or VP1+VP2) of AAV, enabling virus production with only 5 or 10 attachment handles per capsid. Together, our work highlights the importance of site and stoichiometry for generating functional AAV conjugates, and establishes a technology that provides an unprecedented degree of control in manipulating these factors.

## Results and discussion

### Site of attachment profoundly impacts the performance of AAV2-cRGD conjugates

We previously developed a three-plasmid transfection system for packaging AAV serotype 2 (AAV2), with ncAAs (e.g., AzK, Figure 1a) incorporated into its capsid at predefined sites.^*32, 34, 35*^ In this system, the native AAV genes (Rep-Cap) were replaced with a wild-type EGFP reporter in between the inverted terminal repeats (ITRs), and these were supplied *in trans* (Figure 1b, S1). A UAG stop codon was placed in the Cap gene at a site of interest to enable the incorporation of ncAAs. Given the overlapping nature of the three capsid proteins (VP1, VP2, and VP3, present in 5, 5, and 50 copies per capsid, respectively), this results in the incorporation of a ncAA residue in each of the 60 capsid proteins. The archaea-derived pyrrolysyl-tRNA synthetase (PylRS)/tRNA^Pyl^ pair^*26, 28, 40*^ was used to incorporate various ncAAs, including AzK (Figure 1a), in response to the repurposed UAG codon. The incorporation of AzK had minimal impact on AAV infectivity, but it enabled chemoselective labeling of the capsid using strain-promoted azide-alkyne click chemistry (SPAAC). Furthermore, we incorporated AzK at a conserved primary receptor binding site (Arg588) of AAV2 to generate a mutant that is incapable of binding its endogenous receptor (heparan sulfate proteoglycan; HSPG), and labeled it with a cRGD peptide using SPAAC, which retargeted the virus to the αVβ3 integrin receptor.

**Figure 1.**
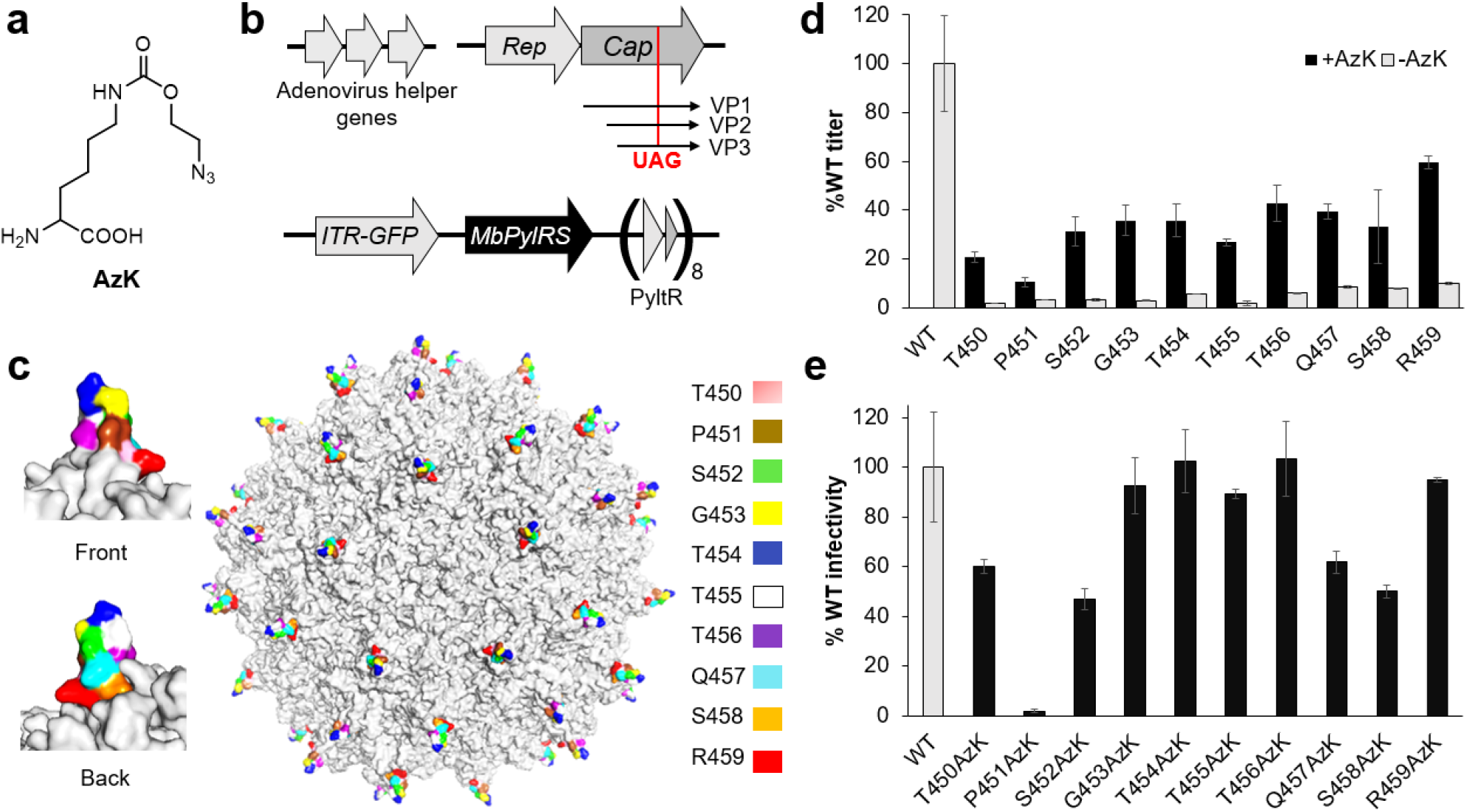
Site-specific incorporation of AzK into the spike region of the AAV capsid. a) Structure of AzK. b) A scheme describing the elements in the plasmid system used to produce ncAA-containing AAV2. c) A color-coded depiction of the distribution of sites targeted for AzK incorporation in the AAV2 capsid. d) Production of various AzK-mutants of AAV2 (packaged genome copies measured by qPCR) in the presence or absence of 1 mM AzK in the media, normalized to the percentage of WT AAV2 titer. e) Infectivity of the AzK-mutants of AAV2, normalized to the percentage infectivity of WT AAV2, measured by the expression of an encoded EGFP reporter, upon infecting HEK293T cells at a constant MOI 50 (also see Figure S3).

This established system to retarget AAV2 through chemical modification provides a great platform to systematically explore how the performance of such conjugates is affected by the labeling site and stoichiometry. To evaluate the impact of the site of attachment, we focused on the highly surface exposed ‘spike’ on the AAV2 capsid, which comprises 10 consecutive amino acid residues (450 to 459; Figure 1c) present in each of the 60 capsid proteins. We individually mutated each of these ten spike residues to a UAG nonsense codon, and used these in our established AAV2 packaging system to create AzK-containing virus. Using qPCR analysis to quantify packaged genome copies, we observed successful AAV2 production for all ten mutants at levels ranging from 10% to 60% relative to wild-type AAV2 (Figure 1d). Consistent with selective incorporation of AzK at the UAG codon, the virus production for all mutants was strongly dependent on the presence of AzK in the growth medium (Figure 1d). We also evaluated the infectivity of each AAV2 mutant (at constant genome copies per cell) toward HEK293T cells, using the expression of the encoded EGFP reporter. Nearly all of the mutants exhibited either comparable or slightly lower infectivity relative to wild-type AAV2 (Figure 1e, S3), with the exception of the P451AzK mutant, which was much less infective. This is not surprising, given the unique properties of proline and its impact on protein structure that may be perturbed upon substitution with AzK.

We have previously demonstrated that to achieve cRGD-mediated retargeting of AAV2 to the integrin receptor, it is essential to first ‘detarget’ the virus from its native primary receptor HSPG, by mutating the relevant residues on the capsid (e.g., R588 and R585).^*32*^ Introducing the integrin-binding cRGD ligand onto a capsid with an intact HSPG binding site results in a sharp loss of infectivity toward cell lines presenting either receptor.^*32*^ In light of this prior observation, we introduced two additional mutations (R585A and R588A; termed AAV2-KO) that abolish HSPG binding in each of the 10 spike UAG mutants described above, such that these could be retargeted through cRGD attachment. We were able to package each of the ten AAV2-KO-AzK spike mutants with good yield (40%-60% of WT-AAV2; Figure S4a), and as expected, these mutants exhibited drastically attenuated infectivity (Figure S4b and S4c) relative to wild-type AAV2. Next, we used SPAAC to attach DBCO-cRGDFC (Figure 2a), a synthetic cRGD peptide connected to an azide-reactive cyclooctyne group, onto each AAV-KO-AzK mutant. The infectivity of the resulting conjugates was evaluated toward SK-OV-3 cancer cell line, which overexpresses the integrin receptor, as well as HEK293T cell line that does not. Attachment of cRGD was found to remarkably increase the infectivity of the AAV2-KO-conjugates toward integrin-expressing SK-OV-3 cells (Figure 2b, S5), while it had negligible impact on their infectivity toward HEK293T cells (Figure S6). Intriguingly, the site of attachment had a dramatic impact on retargeting efficiency. Attachment of cRGD to the central residues (453 - 457) within the spike, which are in general more protruding, resulted in more efficient retargeting; site 456 exhibited the highest retargeting efficiency relative to wild-type AAV2. To further confirm successful retargeting, we demonstrated that the infectivity of the AAV2-KO-456-cRGD toward SK-OV-3 cells can be inhibited by the presence of excess RGD in the media, even though an identical treatment had negligible impact on the wild-type virus (Figure S7). In contrast, the presence of heparin in the media had a limited impact on AAV2-KO-456-cRGD infectivity, even though it strongly inhibited wild-type AAV2 (Figure S7). These studies show that the ncAA mutagenesis technology provides access to a broad array of potential sites to attach function-modulating groups on the AAV capsid, and that the site of attachment has a major impact on the activity of the resulting conjugates.

**Figure 2.**
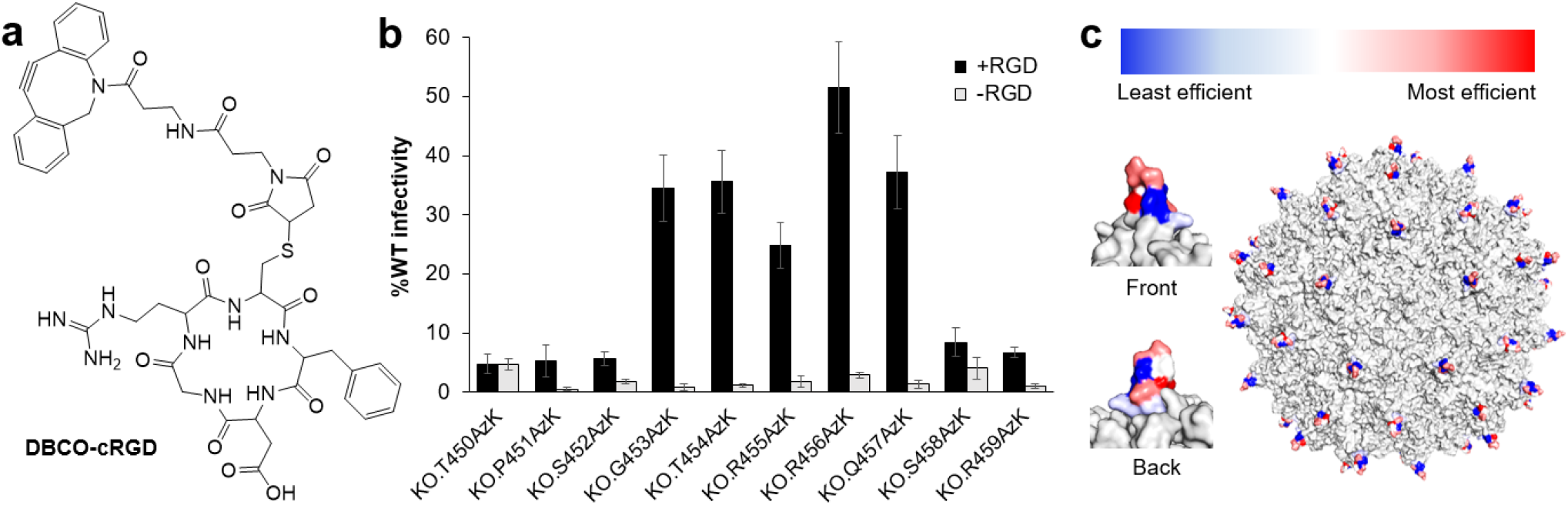
Retargeting efficiency of AAV2-cRGD conjugates is site-dependent. a) Structure of DBCO-cRGD. b) Retargeting efficiency of different AAV2.KO-AzK mutants with and without cRGD-functionalization, shown as a percentage of WT AAV2 infectivity, measured by the expression of an encoded EGFP reporter, upon infecting SK-OV-3 cells at a constant MOI 2500. c) Position of the ten spike residues used in this experiment, color-coded by the efficiency of retargeting.

### Activity of AAV2-cRGD conjugates is sensitive to the stoichiometry of modification

The established platform to generate ncAA-modified AAV produces capsids displaying 60 attachment sites. Although controlled chemical modification of this capsid can produce various degrees of labeling, it is currently unclear how the stoichiometry of capsid modification affects the performance of the corresponding AAV conjugates. To explore this relationship, we sought to create AAV conjugates with increasing numbers of cRGD ligands per capsid through controlled modification. To achieve this, AAV2-KO-T454AzK was incubated with 20 μM DBCO-cRGDFC for increasing lengths of time, followed by quenching the labeling reaction with a large excess of AzK. The infectivity of the resulting conjugates toward integrin-expressing SK-OV-3 cells was evaluated (Figure 3a, S8). As expected, the unmodified AAV2-KO-T454AzK was poorly infective, but upon incubation with DBCO-cRGDFC, its infectivity rapidly increased and reached a peak at 120 min reaction time. However, continued labeling with DBCO-cRGDFC resulted in a sharp decrease in infectivity (Figure 3a, S8), suggesting that an intermediate labeling density is needed for optimal retargeting efficiency. While a low stoichiometry of cRGD labeling is insufficient to drive efficient retargeting, over-modification of the capsid is also detrimental. To estimate the degree of labeling under the optimal conditions, we repeated the capsid labeling of AAV2-KO-T454AzK with 20 μM DBCO-TAMRA for 120 minutes, which enables the visualization of SPAAC-labeling of the capsid through fluorescence imaging (Figure 3b). As a control, we also labeled AAV2-KO-T454AzK with a large excess of DBCO-TAMRA (200 μM) for 24 h, which should result in near-complete modification of all accessible AzK sites on the capsid. A site-specifically TAMRA-labeled reporter protein (GFP) was spiked into AAV2-KO-T454AzK to serve as an internal control for the subsequent imaging analyses. Quantitative imaging analyses comparing the fluorescence intensity of fully TAMRA-labeled AAV2-KO-T454AzK, and the labeling reaction with 20 μM probe quenched at 120 min, revealed approximately 20% labeling (i.e., roughly 12 of the available 60 attachment sites labeled) for the latter (Figure 3b). These observations show that the stoichiometry of labeling is a key determinant of the activity of AAV conjugates. Importantly, the optimal labeling density for retargeting was found to be somewhat low (approximately, 12 per capsid), while the over-modification of the capsid had a significant deleterious effect on its infectivity, possibly due to a perturbation of the delicate capsid functions during entry or downstream trafficking.

**Figure 3.**
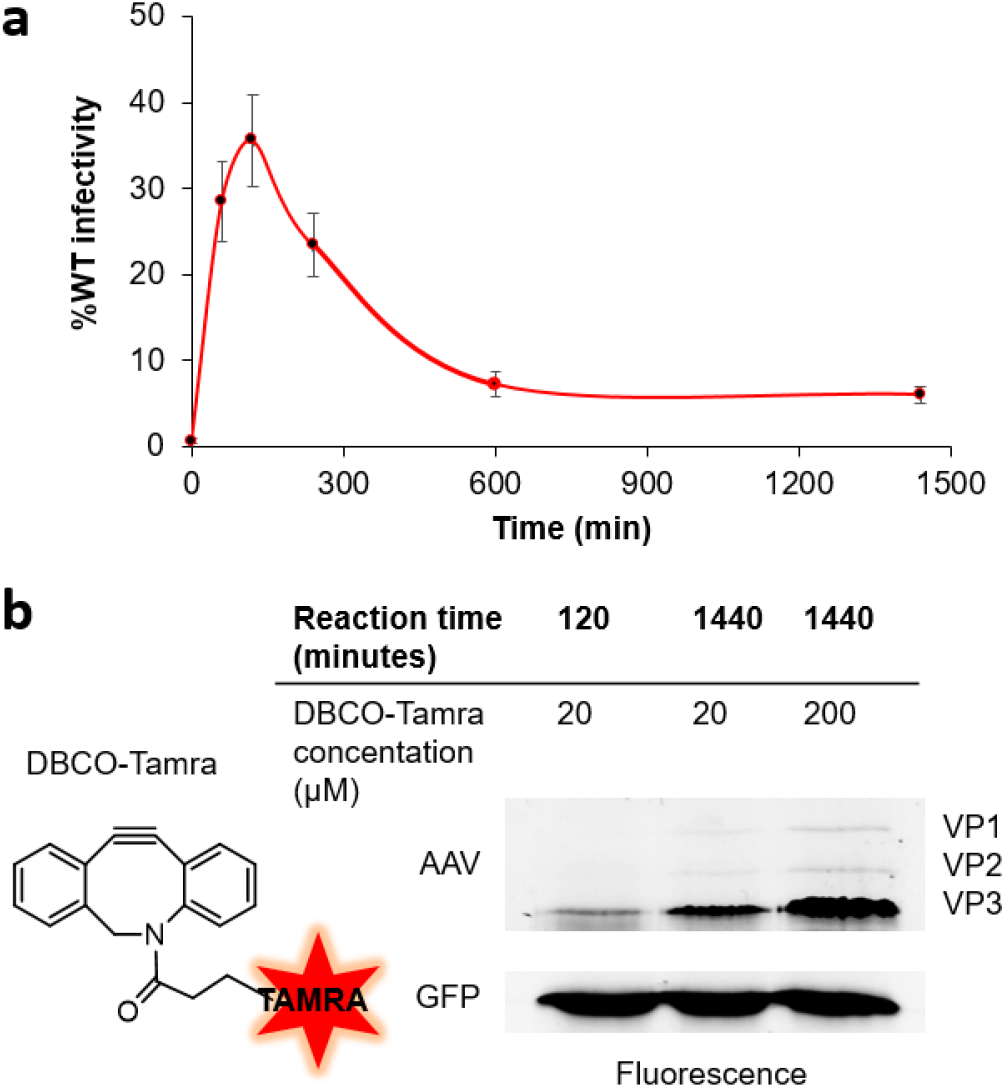
The stoichiometry of capsid labeling affects the retargeting efficiency of AAV-cRGD conjugates. a) Infectivity of cRGD conjugates of AAV2-KO-454AzK, produced by incubating it with 20 μM DBCO-cRGD for different lengths of time (shown in x-axis), shown as a percentage of WT AAV2 infectivity, measured by the expression of an encoded EGFP reporter upon infecting SK-OV-3 cells at a constant MOI of 2500. b) SDS-PAGE followed by fluorescence imaging showing the degree of capsid labeling of AAV2-KO-454AzK upon incubation with 20 μM DBCO-TAMRA for 2 h or 24 h, as well as a large excess of DBCO-TAMRA for 24 h to achieve full labeling. Purified EGFP, pre-labeled with DBCO-TAMRA, was spiked into the virus stock to serve as an internal loading control.

### Controlling the number of attachment sites on AAV through ncAA incorporation into minor capsid proteins

The observations described above highlight the importance of introducing controlled modification onto the AAV capsid, and the perils of over-modification. However, currently, the only way to generate lower stoichiometry of labeling of an AAV capsid harboring 60 ncAAs involves restricting the degree of labeling, which generates a heterogeneous mixture of conjugates with varying labeling stoichiometry. The ability to generate low-stoichiometry conjugates in a homogeneous manner will be valuable, but it would require the ability to introduce only a defined number of ncAA handles per capsid during virus packaging. The AAV capsid is composed of three distinct proteins, VP1, VP2, and VP3, present in 5, 5, and 50 copies per capsid, respectively.^*41*^ The overlapping sequences of these three proteins are encoded in the Cap gene, and are differentially expressed through alternative splicing and start-codon usage. We recognized that the ability to selectively incorporate a ncAA residue in the minor capsid proteins would introduce a lower number of attachment sites per capsid in a defined manner. To achieve this, the expression of the three overlapping capsid proteins must be ‘uncoupled’ from each other (Figure 4a). By introducing mutations in the translation start site(s) of VP1 and/or VP2 (ΔVP1, ΔVP2, or ΔVP1,2), it is possible to selectively abolish the expression of these proteins from Cap. The deleted minor capsid protein(s) can then be supplied back *in trans* from a separate Cap gene, wherein VP3 expression is eliminated by mutating its translation start sites (ΔVP3). We used Western blot analysis to verify that the ΔVP1, ΔVP2, or ΔVP3-Cap constructs are unable to express the relevant capsid proteins when transfected into HEK293T cells (Figure S9), and that the split Cap systems can facilitate the packaging of AAV2 at wild-type titer and infectivity (Figure 4b and 4c).

**Figure 4.**
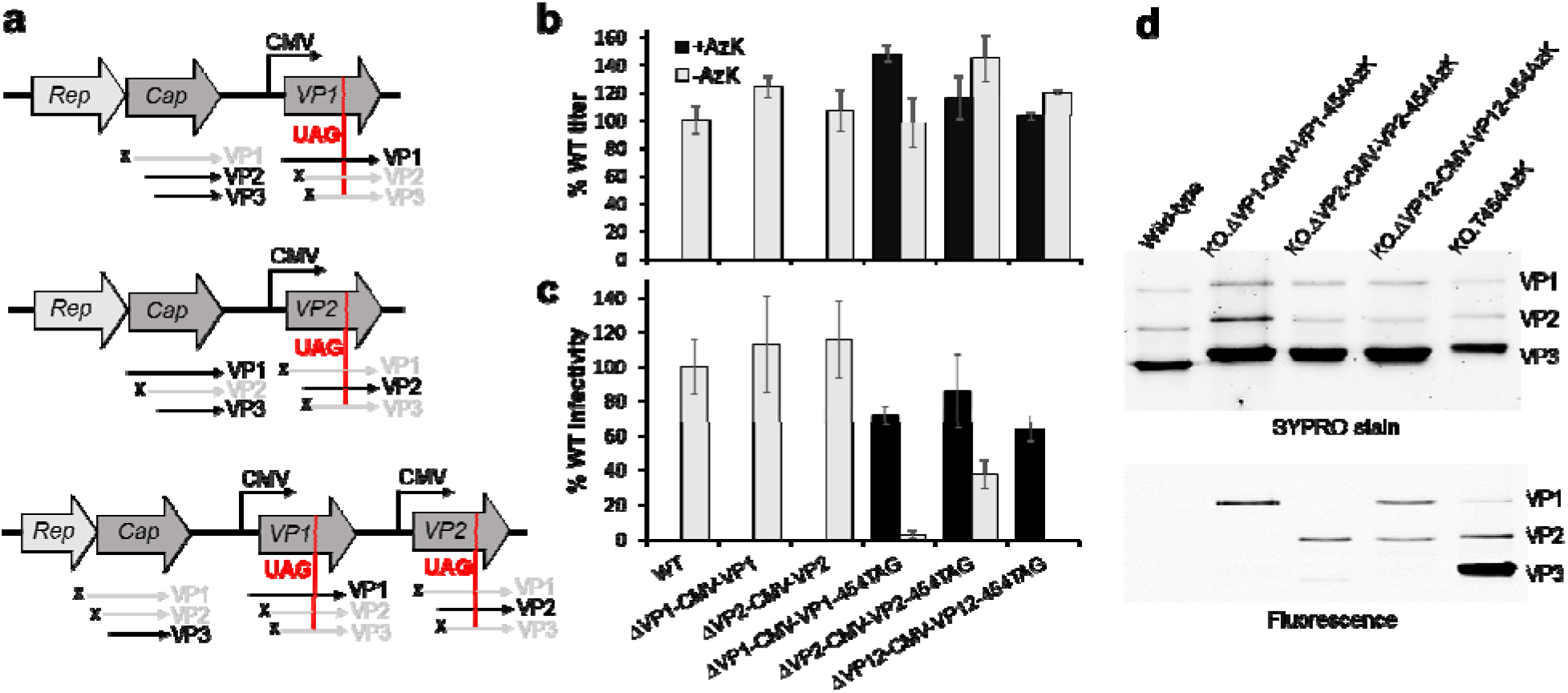
Packaging systems for the selective incorporation of ncAAs into the minor capsid protein of AAV2. a) A scheme describing the strategy to uncouple the expression of VP1 and VP2 from the Cap gene, and expressing these separately from CMV, which enables selective incorporation of AzK into desired minor capsid proteins. b) Using the split-Cap plasmid system, production of WT AAV2, or mutants incorporating AzK at specific minor capsid proteins, measured as packaged genome copies (qPCR) and normalized relative to WT titer. c) Infectivity of the virus preparations (WT AAV2 and AzK-mutants at specific minor capsid proteins) described in panel b, measured by the expression of an encoded EGFP reporter upon infecting HEK293T cells at a constant MOI of 50. d) Incubating these capsids with DBCO-TAMRA, followed by SDS-PAGE and fluorescence imaging confirms our ability to selectively label a distinct subset of the capsid proteins.

Next, in this split-Cap packaging system, we incorporated a UAG codon at the T454 position of the CMV-driven VP1 and VP2 genes, which should facilitate ncAA incorporation only at these minor capsid proteins. Using this strategy, viruses encoding AzK at the T454 position of VP1, or VP2, or both minor capsid proteins were produced at near wild-type levels (Figure 4b) and with comparable infectivity (Figure 4c). It should be noted that high levels of packaged genomes were observed by qPCR both in the presence or absence of AzK (Figure 4b), because even in the absence of minor capsid protein expression, VP3 alone is known to encapsulate the genome.^*42, 43*^ However, such capsids devoid of VP1 showed severely attenuated infectivity (Figure 4c, S10). Labeling the resulting purified AAV2 with DBCO-TAMRA, and subsequent analysis by SDS-PAGE followed by fluorescence imaging confirmed the incorporation of AzK at intended minor capsid proteins (Figure 4d).

### Defined AAV2-cRGD conjugated with a low stoichiometry of labeling

The ability to selectively incorporate AzK at VP1, or VP2 or VP1+VP2, introduces approximately 5, 5, and 10 attachment handles per capsid. To evaluate the retargeting behavior of AAV2-cRGD conjugates produced from these capsids, we labeled them with 20 μM DBCO-cRGD for different lengths of time to generate conjugates with increasing stoichiometry, and analyzed their infectivity toward SK-OV-3 cells, similar to the experiment described in Figure 3a. It should be noted that the capsids used for the retargeting experiment also had mutations R585A and R588A (KO) to detarget them from the native HSPG receptor. Unlike AAV2-KO-454-AzK (60 attachment handles per capsid), which first gained and then sharply lost infectivity with increasing degree of cRGD labeling, the capsids with 5 or 10 attachment handles steadily gained infectivity and reached a plateau (Figure 5a, S11). This suggests that up to 10 cRGD modifications per capsid are tolerated reasonably well without significant adverse effect on infectivity. However, maximal retargeting efficiency of conjugates derived from capsids with 5 AzK residues (either on VP1 or VP2 alone) was significantly lower relative to the capsid with 10 AzK residues (both on VP1 and V2). This is consistent with our previous assessment that optimal retargeting efficiency is achieved at approximately 12 cRGD residues per capsid (Figure 3). We also evaluated the impact of introducing a long and flexible PEG linker between the capsid and the cRGD, which did not significantly alter the retargeting behavior (Figure 5b; S12).

**Figure 5.**
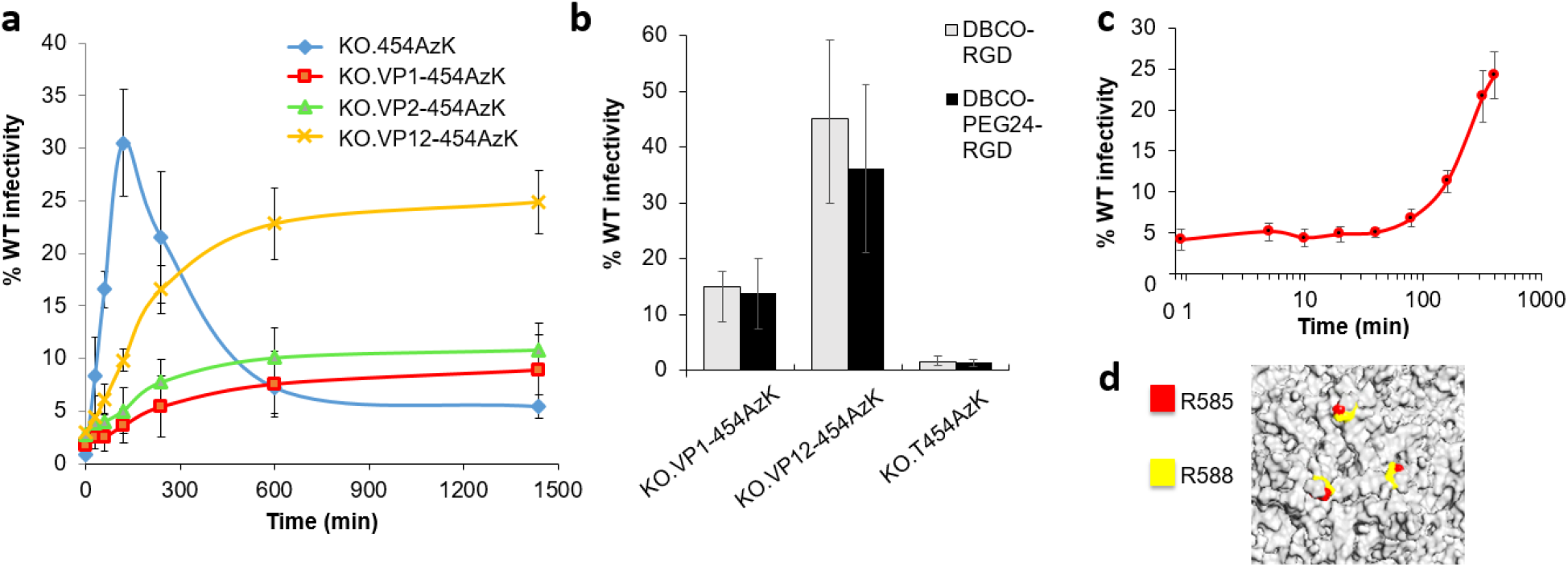
Retargeting behavior of AAV capsids containing a defined number of attachment sites. a) Infectivity of cRGD conjugates of AAV2-KO-454AzK mutants with the AzK in all capsid proteins (60 copies per capsid), in only VP1 or VP2 (5 copies), or in both VP1 and VP2 (10 copies), produced by incubating it with 20 μM DBCO-cRGD for different lengths of time (shown in x-axis), shown as a percentage of WT AAV2 infectivity, measured by the expression of an encoded EGFP reporter upon infecting SK-OV-3 cells at a constant MOI of 2500. b) AAV-cRGD conjugates generated using 50 μM DBCO-cRGD for 24 h with or without a long flexible linker exhibit comparable retargeting efficiencies. c) Incubation of AAV2-KO-VP1,2-454-AzK with 20 μM cRGD-DBCO results in an increase in infectivity toward SK-OV-3 cells, but only after a significant lag period. The infectivity was measured by the expression of an encoded EGFP reporter upon infecting SK-OV-3 cells at a constant MOI of 2500, and shown as a percentage of WT AAV2 infectivity. d) Magnified image of the HSPG binding site on AAV2 capsid, with key residues R585 and R588 shown in red and yellow, shows three individual binding sites arranged in a tight cluster around a three-fold axis of symmetry.

The improved efficiency of the AAV2 conjugate with 10 cRGD per capsid relative to that with 5 is indicative of a need for a multivalent binding for efficient retargeting, where multiple cRGD ligands simultaneously associate with their receptors. This is further supported by the observation that when AAV2-KO-VP1,2-AzK is incubated with 20 μM DBCO-cRGD, its infectivity toward SK-OV-3 cells does not start to increase during the first hour of incubation (Figure 5c; S13). This significant ‘lag-period’ indicates that the low-stoichiometry conjugates generated during the early phase of the reaction are poorly infective, and only when more densely labeled conjugates are produced upon further incubation, the retargeting efficiency starts to increase. This is not surprising, given the relatively modest monovalent binding affinity of cRGD peptides,^*44*^ which may not be sufficient to drive efficient capsid-receptor interaction. It is also reminiscent of the multivalent nature of native primary receptor binding by AAV2, where three binding sites are tightly clustered around a three-fold axis of symmetry (Figure 5d). It is conceivable that the use of retargeting ligands with higher binding affinity may enable optimal retargeting at a lower stoichiometry of labeling.

## Conclusions

Precise chemical modification of the capsid offers much potential to introduce therapeutically attractive features in existing AAV vectors for gene therapy. Here we show that the ncAA mutagenesis technology provides an outstanding strategy to this end. Small ncAA residues with bioorthogonal conjugation handles are well tolerated across numerous exposed sites on the capsid, providing countless ways to introduce selective chemical modifications at predefined sites. We demonstrate that the attachment site plays a crucial role in the performance of the AAV2 conjugates, highlighting the importance of having the flexibility to select the conjugation site from a large catalog of potential options. Additionally, we reveal how the stoichiometry of capsid labeling is a crucial factor that determines the activity of AAV conjugates. In particular, over-modification of the capsid was found to be detrimental. To enable the synthesis of defined AAV conjugates with lower labeling stoichiometry, we developed technology to selectively introduce ncAA handles into the minor capsid proteins of AAV, which introduces 5 or 10 attachment handles per capsid. Our work establishes a platform to modify the AAV capsid with an unprecedented degree of control over site and stoichiometry. Together, it enables systematic optimization of the performance of functional AAV conjugates, akin to structure-activity relationship strategies used in medicinal chemistry for fine-tuning small-molecule therapeutics.

## ASSOCIATED CONTENT

### Supporting Information

Experimental methods, nucleotide sequences, supplementary figures and tables. (PDF)

## AUTHOR INFORMATION

### Notes

A patent application on precise capsid modification technology has been submitted. AC is a cofounder and senior advisor of BrickBio, Inc.

## ACKNOWLEDGMENT

This work was supported by NIH (R35GM136437) and NSF (MCB 1817893) to A.C.

