## Supporting information for "Precise manipulation of site and stoichiometry of capsid modification enables optimization of functional adeno-associated virus conjugates"

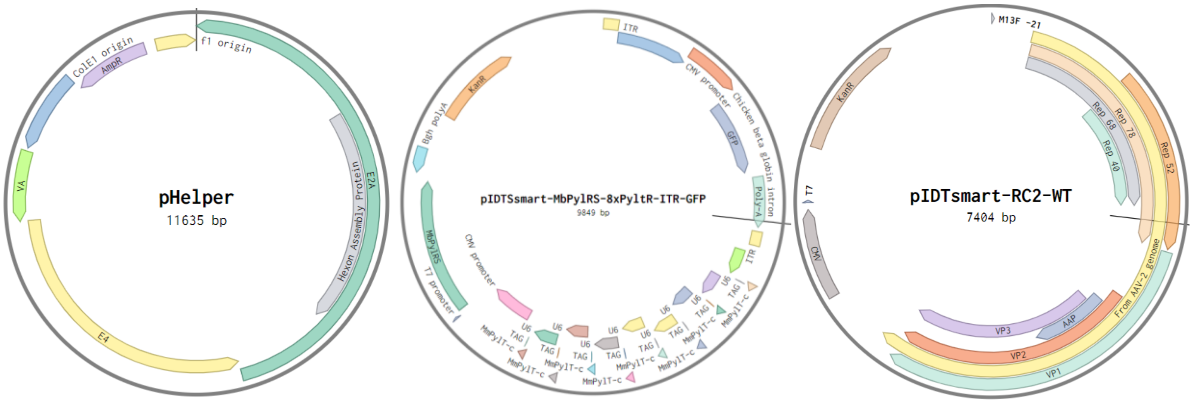


| Construct | Deleted start codons in native Cap | Deleted start codons in CMV-Cap |
|---|---|---|
| pIDTsmart-RC2-ΔVP1-CMV-VP1 | M1L (ATG→CTC) | T138T(ACG→ACC),<br>T197T(ACG→ACC),<br>L202L(CTG→CTC),<br>M203L(ATG→CTC),<br>M211L(ATG→CTC),<br>M235L(ATG→CTC) |
| pIDTsmart-RC2-ΔVP2-CMV-VP2 | T138T(ACG→ACC) | L202L(CTG→CTC),<br>M203L(ATG→CTC),<br>M211L(ATG→CTC),<br>M235L(ATG→CTC) |
| pIDTsmart-RC2-ΔVP1,2-CMV-VP1,2 | M1L (ATG→CTC),<br>T138T(ACG→ACC) | <u>CMV-VP1</u>: T138T(ACG→ACC),<br>T197T(ACG→ACC),<br>L202L(CTG→CTC),<br>M203L(ATG→CTC),<br>M211L(ATG→CTC),<br>M235L(ATG→CTC);<br><u>CMV-VP2:</u> T197T(ACG→ACC),<br>L202L(CTG→CTC),<br>M203L(ATG→CTC),<br>M211L(ATG→CTC),<br>M235L(ATG→CTC) |

**Figure S1**. Plasmids used for producing ncAA containing AAVs. pHelper contains the adenoviral E2A, E4, and VA genes. pIDTSmart-MbPylRS-8xPytR-ITR-GFP contains wild-type *M. barkeri* pyrrolysyl synthetase driven by a CMV promotor, eight copies of the *M. mazei* pyrrolysyl $tRNA_{CUA}$ expression cassette driven by a human U6 promoter, and a CMV-EGFP cargo flanked by packaging signals (AAV2 ITRs). pIDTSmart-RC2-WT contains the *cap* and *rep* gene from the AAV2 genome. TAG mutations at specific sites of the *cap* gene define the site of ncAA incorporation. For retargeting viruses, HSPG binding sites are mutated R585A(AGA→GCC), R588A(AGA→GCC). The table lists mutations introduced to abolish expression of different capsid proteins

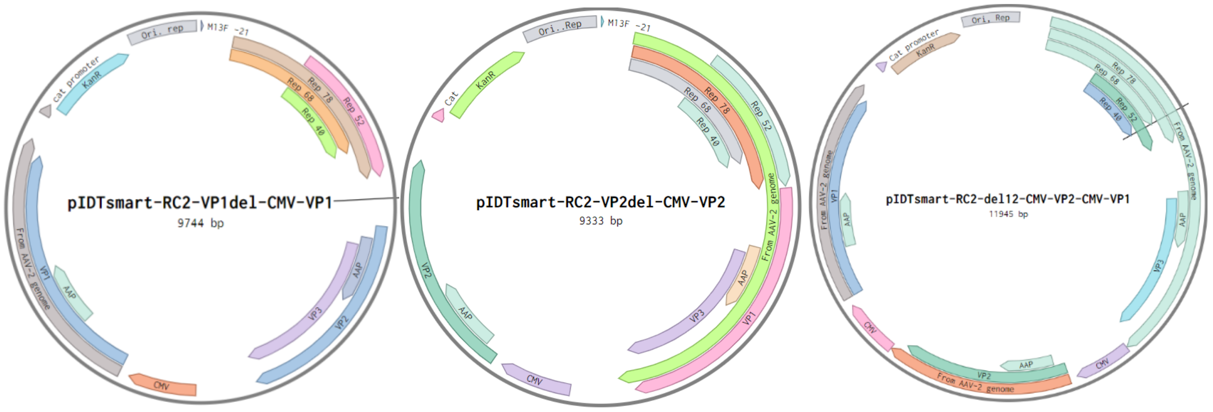


**Figure S2**. Plasmids used for producing AAVs with ncAA incorporated at selected capsid proteins. Selected capsid proteins are uncoupled from AAV2 genome by deleting their start codons. Single capsid protein genes are provided back by deleting major capsid protein VP3's and other other minor capsid protein's start codons, driven by a CMV promoter. TAG mutations at specific sites of the *cap* gene define the site of ncAA incorporation.

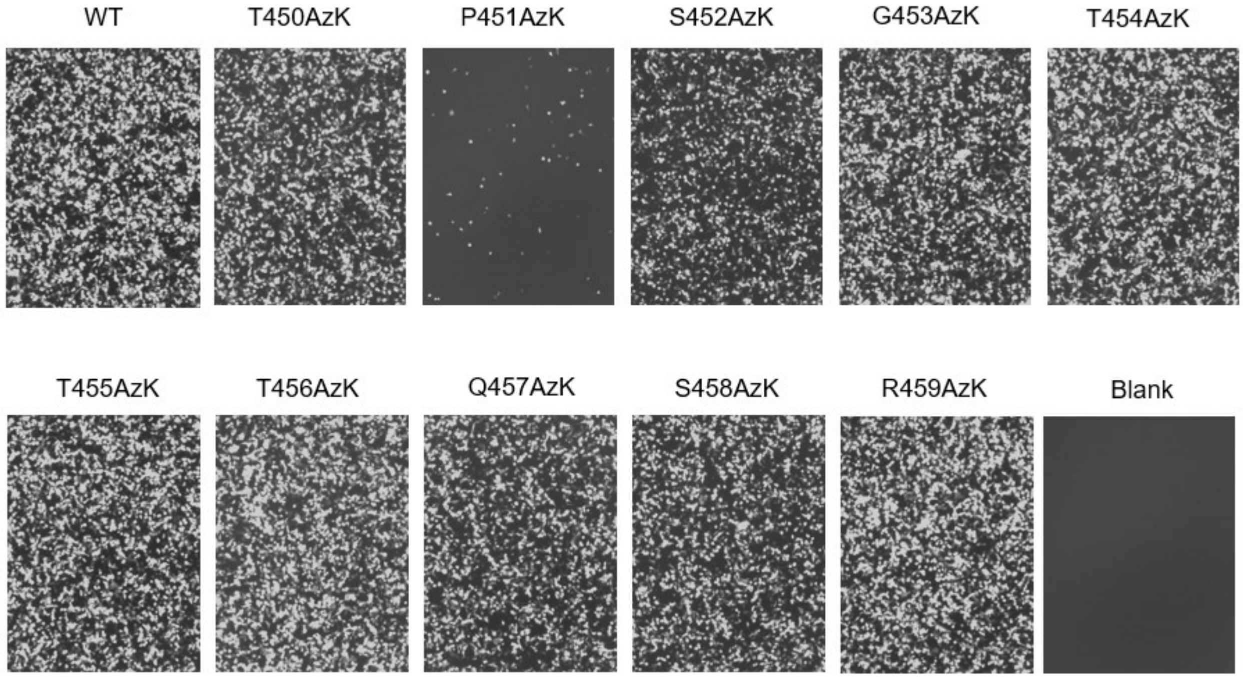


**Figure S3**. Fluorescence images of cells associated with the experiment described in Figure 1e. HEK293T cells were infected with a constant MOI (50) of wild-type or AzK mutants of AAV2, and imaged 48 h post-infection.

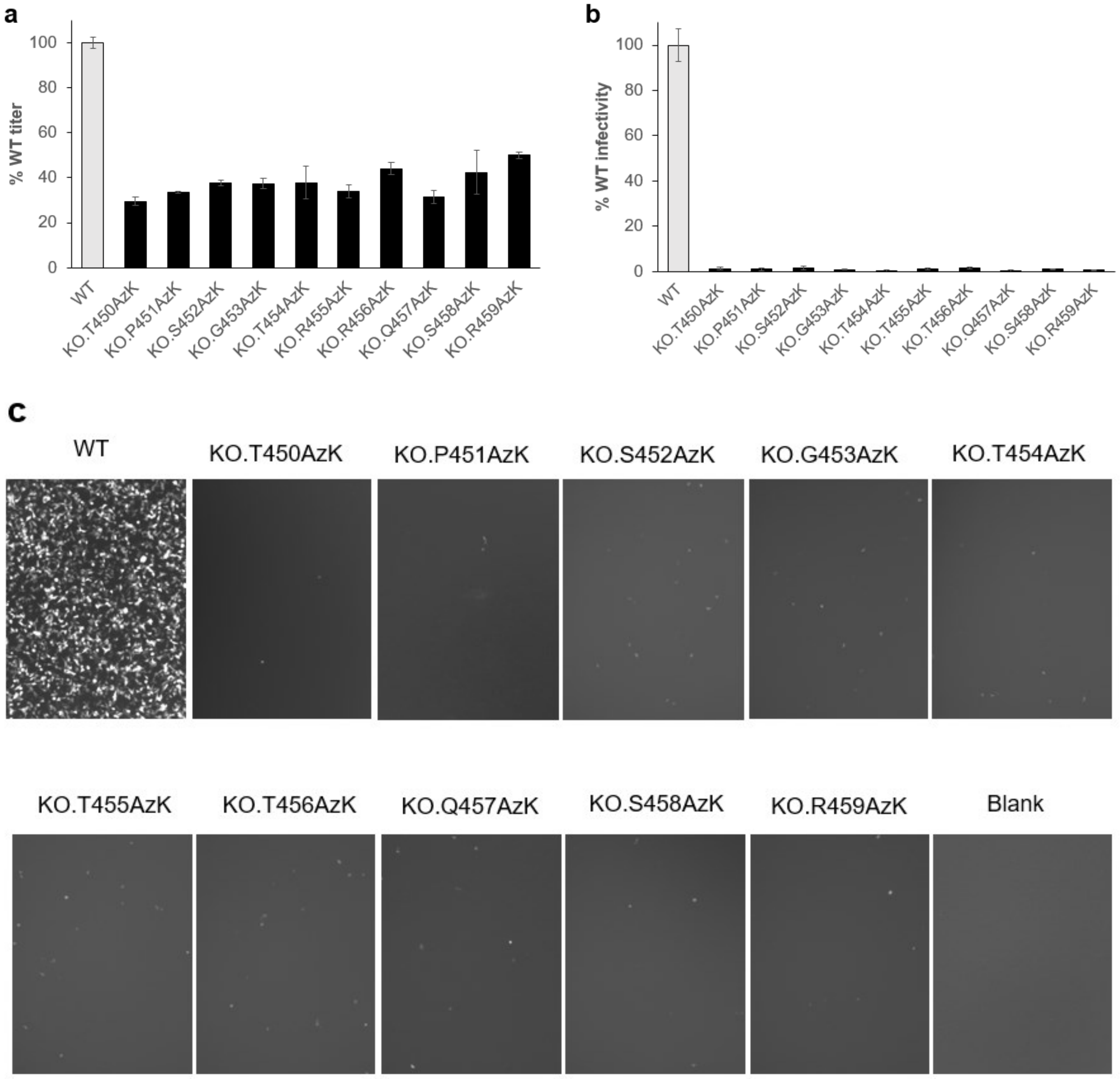


**Figure S4**. a) Production of various AzK-mutants of AAV2-KO (packaged genome copies measured by qPCR) normalized to the percentage of WT AAV2 titer. b) Infectivity of the AzK-mutants of AAV2, normalized to the percentage infectivity of WT AAV2, measured by the expression of an encoded EGFP reporter, upon infecting HEK293T cells at a constant MOI 50. c) EGFP-fluorescence images of cells associated with the experiment described in panel b. HEK293T cells were infected with a constant MOI (50) of wild-type or site-specific AzK mutants of AAV2-KO, and imaged 48 h post-infection.

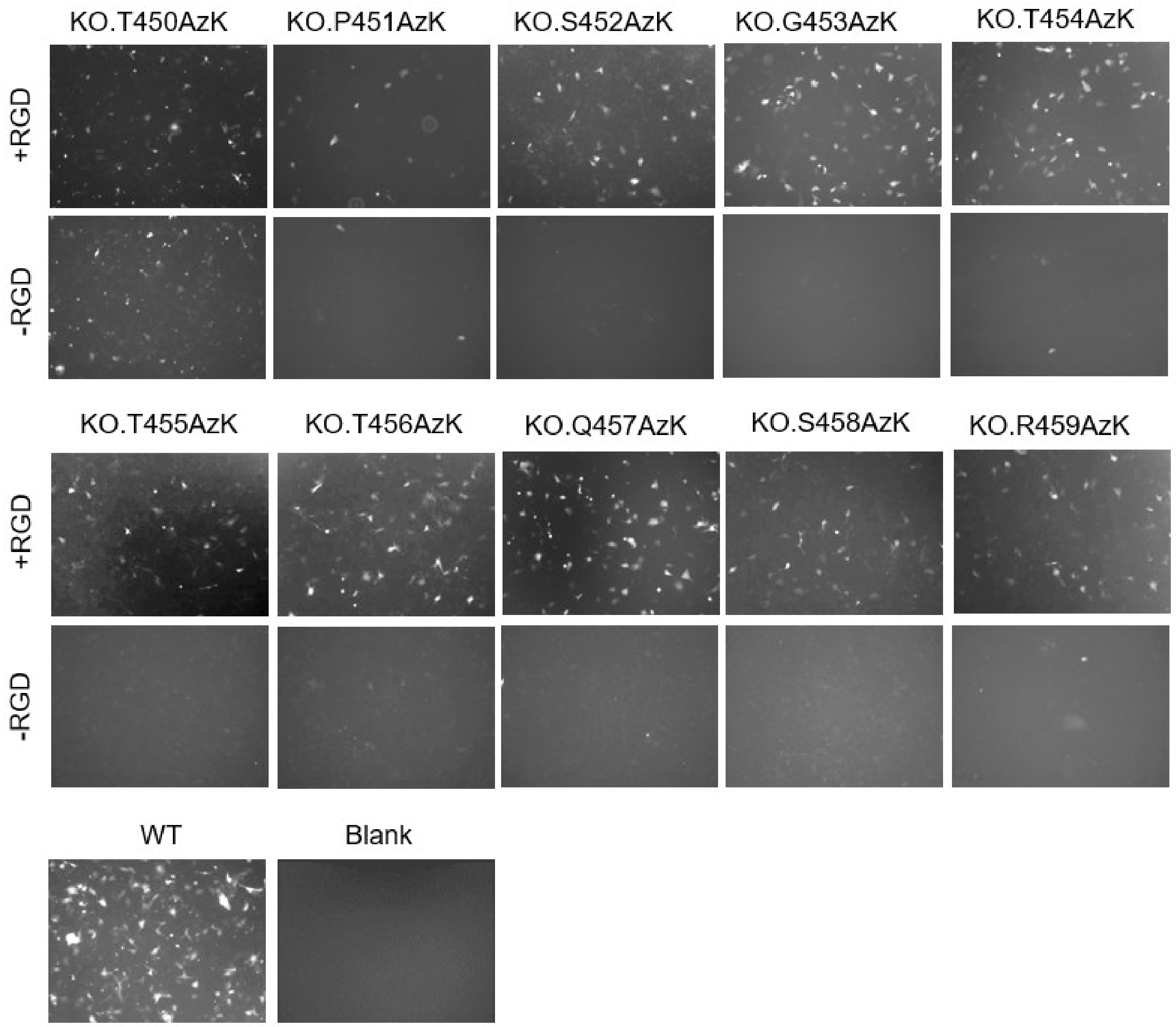


**Figure S5**. Fluorescence images of cells associated with the experiment described in Figure 2b. SK-OV-3 cells were infected with a constant MOI (2500) of wild-type or site-specific AzK mutants of AAV2-KO with or without reacting with 20 μM DBCO-cRGDFC for 2 h, and imaged 48 h post-infection.

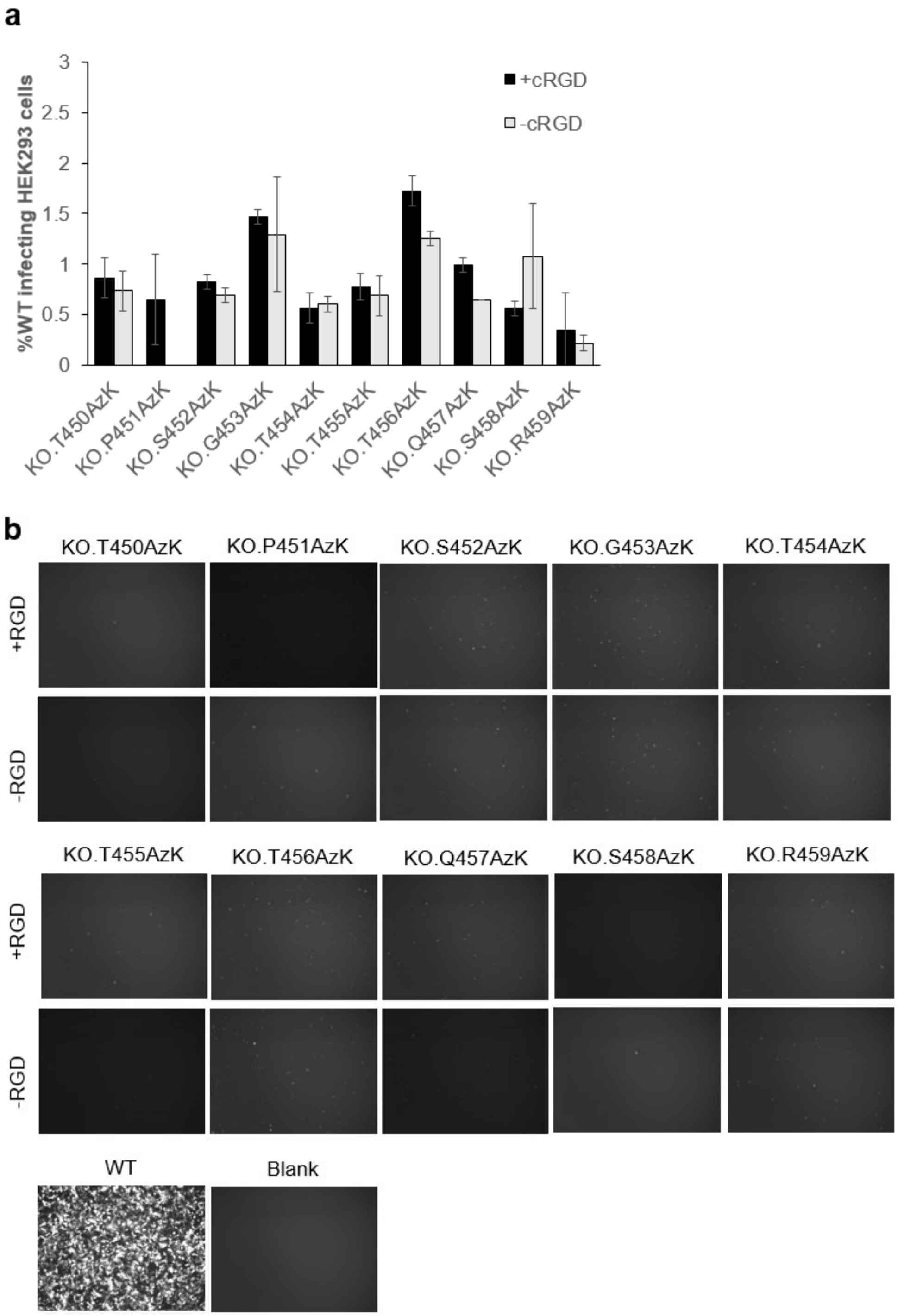


**Figure S6**. a) Various detargeted AAV2.KO-AzK mutants show low infectivity toward integrin-deficient HEK293 cells, with or without cRGD labeling (infectivity shown relative to WT AAV2). b) EGFP-fluorescence images associated with the experiment described in Figure S6a. HEK293T cells were infected with a constant MOI (50) of WT-AAV2 or different AzK mutants of AAV2-KO with or without labeling with 20 μM DBCO-cRGD for 2 h, and imaged 48 h post-infection.

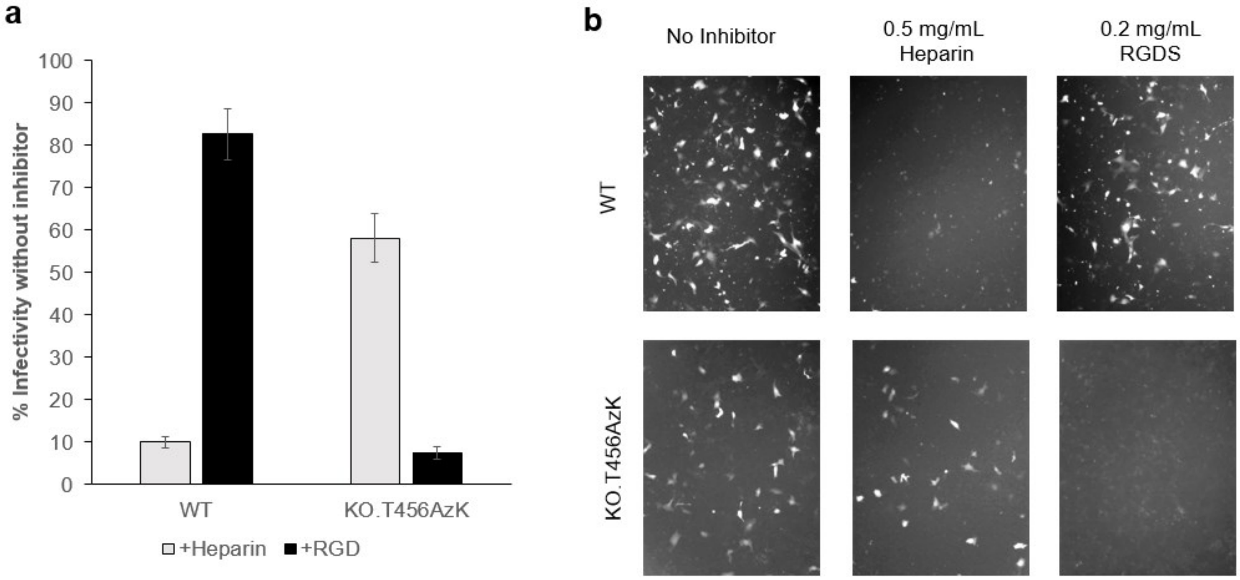


**Figure S7**. a) Infectivity of wild-type AAV2 and AAV2-KO-T456AzK functionalized with cRGDFC towards SK-OV-3 cells in the presence of free RGD peptide or heparin. Relative infectivity was measured as the percentage of EGFP expressing cells (FACS), and was normalized relative to the corresponding no inhibitor control. b) Fluorescence images of cells associated with panel a.

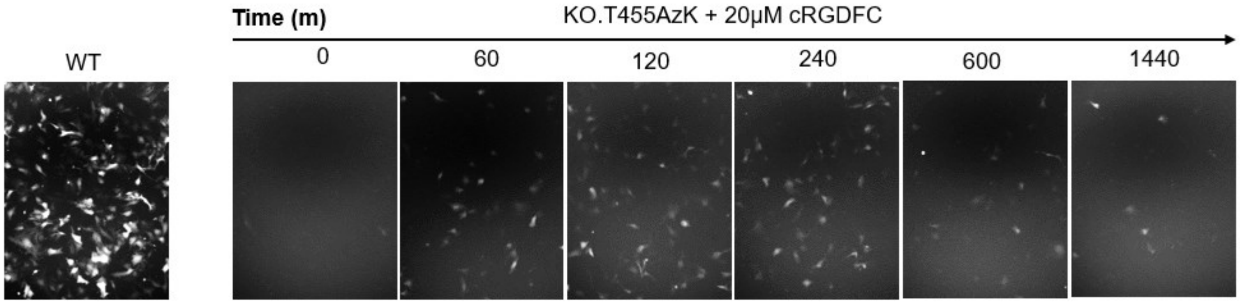


**Figure S8**. Fluorescence images of cells associated with the experiment described in Figure 3a. SK-OV-3 cells were infected with a constant MOI (2500) of wild-type AAV2, or AAV2-KO-T454AzK incubated with 20 µM DBCO-cRGDFC for indicated periods of time, and imaged 48 h post-infection.

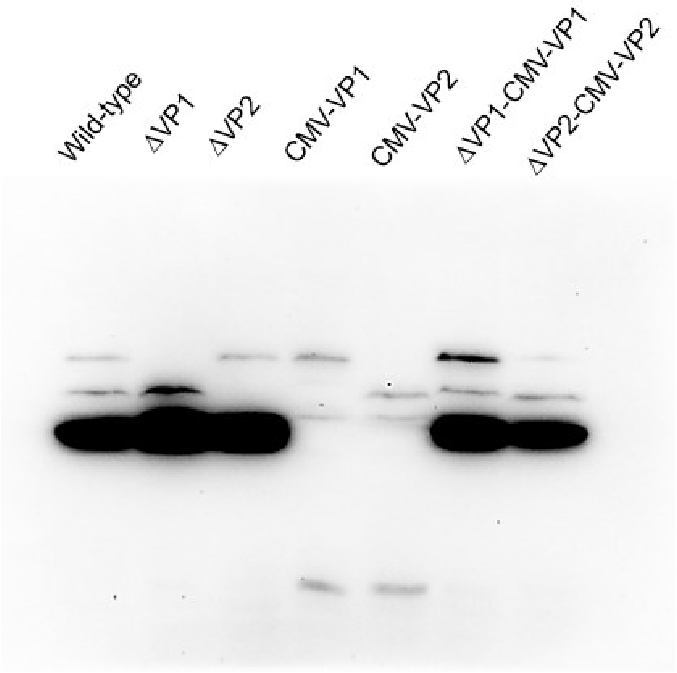


**Figure S9**. Western blot analysis of AAV2 capsid proteins expression from different Cap-constructs upon transiently transfecting them into HEK203T cells. Equal volumes of lysates were resolved by 10% SDS-PAGE, and analyzed by Western blot using anti-AAV B1 antibody. ∆VP1 and ∆VP2 constructs represent constructs where translation start sites of VP1, or VP2, respectively, were mutated to abolish their expression. CMV-VP1 and CMV-VP2 represent constructs designed to only express VP1 and VP2, respectively, but not VP3. Very low levels of VP3 expression from CMV-VP1 and CMV-VP2 are still observed, despite extensive translation start site engineering (Figure S1).

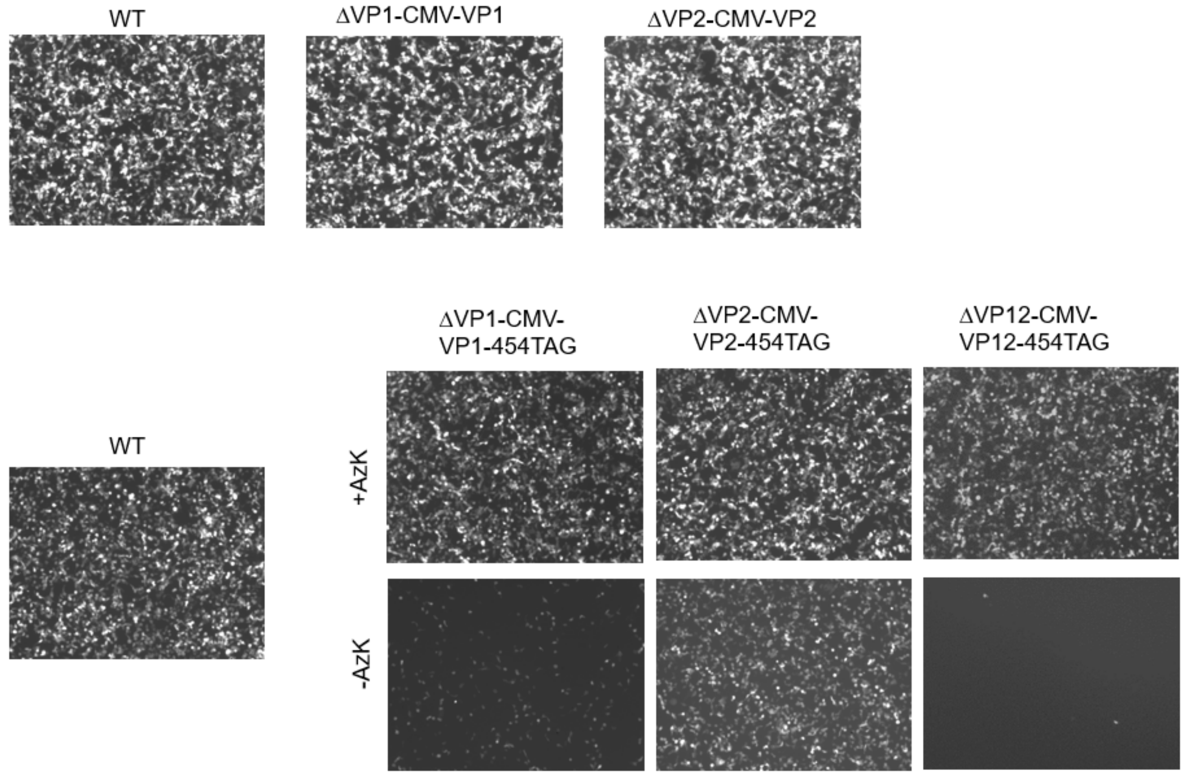


**Figure S10**. EGFP-fluorescence images associated with the experiment described in Figure 4c. HEK293T cells were infected with a constant MOI (50) of wild-type, or site-specific AzK mutants at selected capsid proteins of AAV2, and imaged 48 hr post-infection.

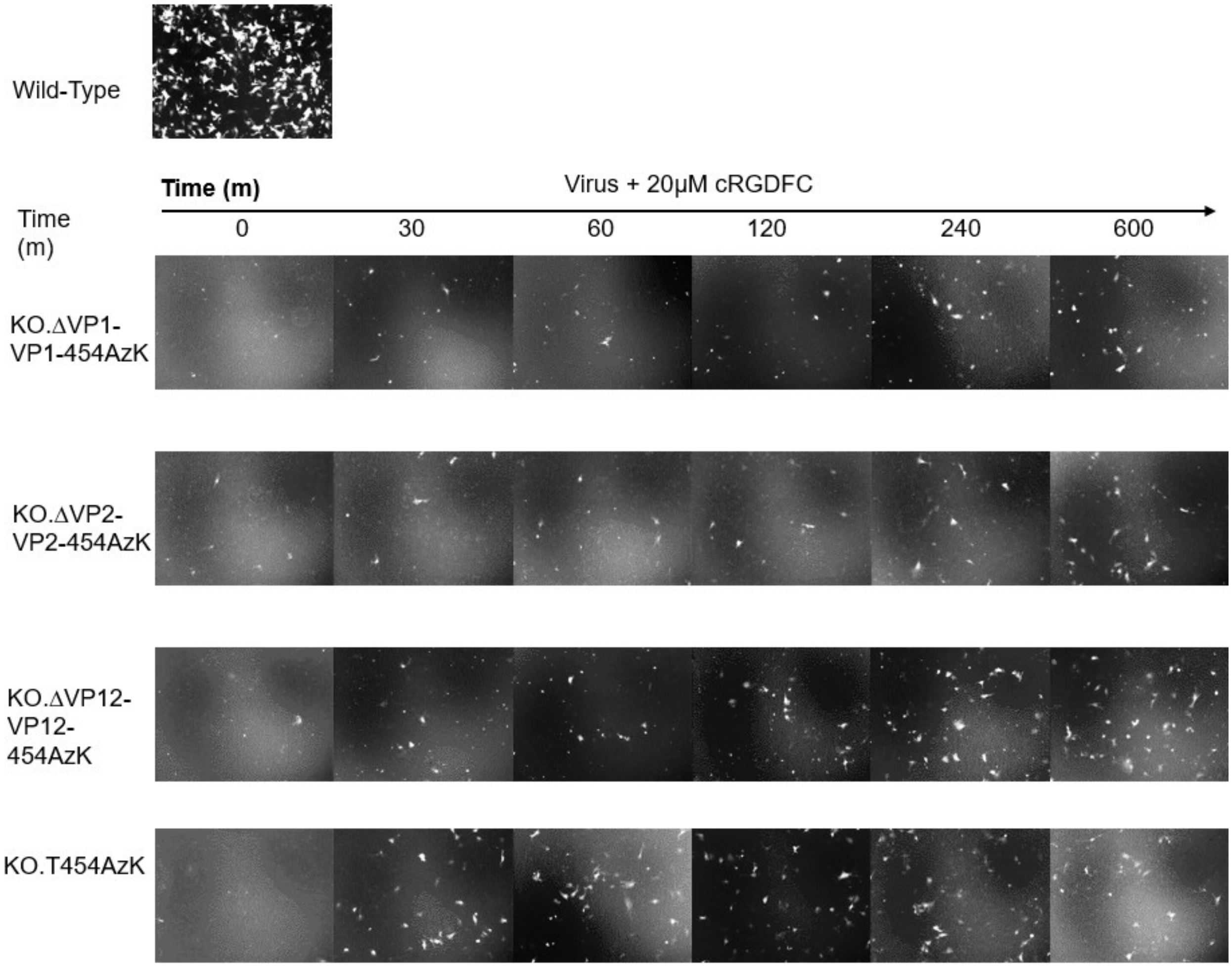


**Figure S11**. Fluorescence images of cells associated with the experiment described in Figure 5a. SK-OV-3 cells were infected at a constant MOI (2500) of wild-type AAV2, or AAV2-KO-AzK mutants at specified capsid proteins upon reacting with 20 μM DBCO-cRGDFC for specified lengths of time, and imaged 48 h post-infection.

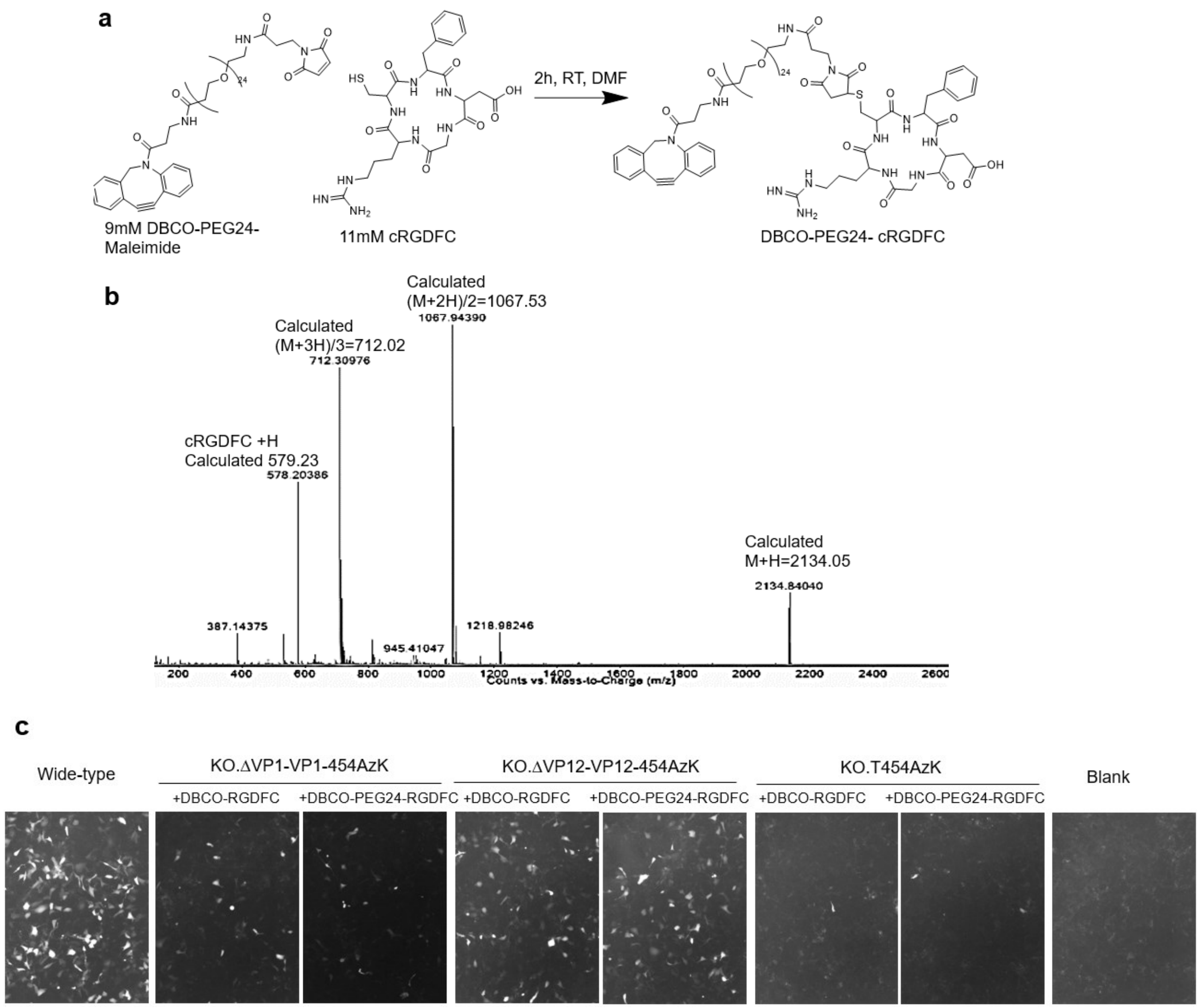


**Figure S12**. a) Synthesis scheme of DBCO-PEG24-cRGDFC. b) MS spectrum of DBCO-PEG24-cRGDFC. c) EGFP-fluorescence images associated with the experiment described in Figure 5b. SK-OV-3 cells were infected with a constant MOI (2500) of wild-type or T454AzK at selected capsid proteins of AAV2-KO reacting with 50 μM DBCO-cRGDFC for 24 hours, and imaged 48 h post-infection.

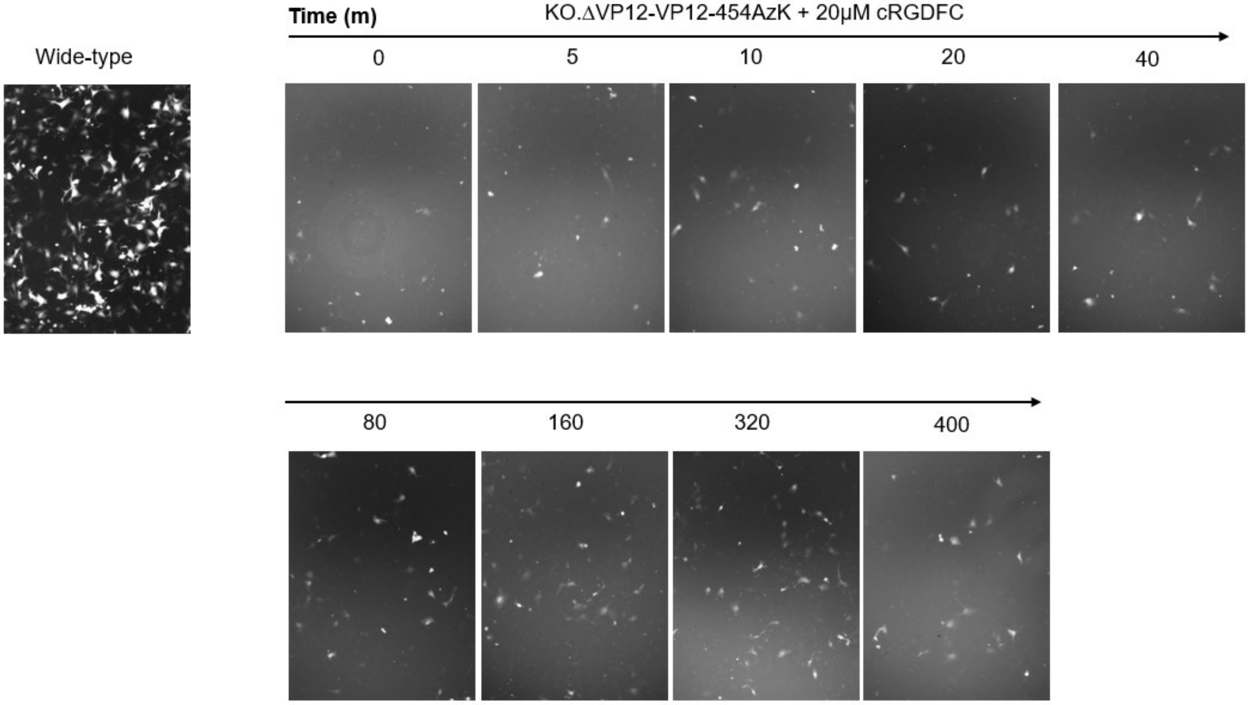


**Figure S13**. EGFP-fluorescence images associated with the experiment described in Figure 5c. SK-OV-3 cells were infected with a constant MOI (2500) of AAV2-KO-VP1,2- 454AzK after reacting with 20 µM DBCO-cRGDFC for indicated lengths of time, and imaged 48 h post-transfection.

### Materials and Methods

**Cell Culture.** HEK293T cells and SK-OV-3 cells were obtained and maintained as previously described.[1]

**Cloning and Plasmids.**

Transformation was done in Top10 cells using BioRad electroporator. DNA oligo synthesis and Sanger Sequencing were performed by Genewiz. Phusion polymerase was purchased from Thermo Scientific™, PrimeSTAR® Max DNA Polymerase was purchased from Takara, T4 DNA Ligase was purchased from Qiagen.

pIDTSmart-*Mb*PylRS-8xPyltR-ITR-GFP was generated by inserting 8xU6-*Mm*PyltR cassette at SpeI site and ITR-EGFP-ITR at SbfI site of the plasmid pIDTsmart-*Mb*PylRS.[1] The two inserts 8xPyltR cassette and ITR-EGFP-ITR were digested from the previously described plasmid pIDTSmart-8xPyltR-ITR-EGFP-ITR[1] with restriction enzymes AvrII+NheI and SbfI, respectively.

pIDTSmart-RC2-WT was made by digesting pIDTSmart-RC2-*Mb*PylRS[1] with AvrII and NheI and ligating the plasmid back together, removing the *Mb*PylRS coding region. For the incorporation of ncAA into 60 AAV capsid proteins, spike residue from 450 to 459 was individually replaced by TAG codon by QuickChange site-directed mutagenesis from pIDTSmart-RC2-WT.

VP1 was uncoupled from RC2 genome by mutating its start codon M1L (TAG→CTC) by overlap-extension PCR using VP1-M1L-F, VP1-M1L-R, RC2-HindIII-F, Cap2-SbfI-R primers. VP2 was uncoupled from RC2 genome by mutating its start codon T138T (ACG →ACC) by overlap-extension PCR using VP2-T138T-F, VP2-T138T-R, RC2-HindIII-F, Cap2-SbfI-R primers. A total of five mutations were made to fully abolish VP3 expression. VP3 start codon M235L (ATG→CTC) was mutated by QuickChange site-directed mutagenesis from pIDTSmart-RC2-WT using VP3-M235L-F and VP3-M235L-R primers. Using the resulting plasmid as a template, two other VP3 start codons M203L (ATG→CTC), M211L (ATG→CTC) and two additional alternative starts T197L (ACG→ACC), L202L (CTG→CTC) were mutated by overlap-extension PCR using VP3-T197L/L202L/M203L/211L-F, VP3-L202L/M203L/211L-R, RC2-HindIII-F, Cap2-SbfI-R primers. The overlap PCR products were digested and cloned into pIDTSmart-RC2-CMV-MbPylRS using HindIII and SbfI restriction sites, yielding the plasmid pIDTSmart-RC2-ΔVP1-CMV-MbPylRS, pIDTSmart-RC2-ΔVP2-CMV-MbPylRS, pIDTSmart-RC2-ΔVP3-CMV-MbPylRS.

VP1 was expressed separately from the cap gene by abolishing the expression of VP2 and VP3. This was done by overlap-extension PCR using VP2-T138T-F, VP2-T138T-R, VP1-NheI-F, Cap2-AvrII-R primers on pIDTSmart-RC2-ΔVP3-CMV-MbPylRS as the template. VP2 was expressed separately from the cap gene by removing the VP1-N terminus sequence and expression of VP3, while T138 was mutated to ATG, a stronger start codon for VP2 expression. This was done by amplifying pIDTSmart-RC2-ΔVP3-CMV-MbPylRS using VP2-ATG-NheI-F and Cap2-AvrII-R primers. The PCR products were digested and cloned into pIDTSmart-RC2-ΔVP1-CMV-MbPylRS, pIDTSmart-RC2-ΔVP2-CMV-MbPylRS using NheI and AvrII sites, yielding pIDTSmart-RC2-ΔVP1-CMV-VP1 and pIDTSmart-RC2-ΔVP2-CMV-VP2, respectively.

VP1 and VP2 were uncoupled from RC2 genome by overlap-extension PCR using primers VP2-T138T-F, VP2-T138T, RC2-HindIII-F, Cap2-SbfI-R on pIDTSmart-RC2-ΔVP1-CMV-

MbPylRS as a template. The overlap PCR product was digested and cloned into pIDTSmart-RC2-ΔVP1-CMV-VP1 using HindIII and SbfI restriction sites, yielding the intermediate plasmid pIDTSmart-RC2-ΔVP1,2-CMV-VP1. CMV-VP2 was amplified from pIDTSmart-RC2-ΔVP2-CMV-VP2 using SbfI-CMV-F and Cap2-MluI-R. The PCR product was digested and cloned into pIDTSmart-RC2-ΔVP12-CMV-VP1 using SbfI and MluI sites to yield pIDTSmart-RC2-ΔVP1,2-CMV-VP1,2.

From these plasmids, stop codons were incorporated into the final plasmids by performing overlap-extension PCR on appropriate template and ligated into the corresponding plasmid for the incorporation of ncAA.

**Primers used in this study**:

| Primer name | Sequence |
|---|---|
| T450TAG-F | GAGCAGAACAAACTAGCCAAGTGGAACCACCACGCAGTCAAG |
| T450TAG-R | GTGGTTCCACTTGGCTAGTTTGTTCTGCTCAAGTAATACAGGTACTGGTC |
| T451TAG-F | GAGCAGAACAAACACTTAGAGTGGAACCACCACGCAGTCAAG |
| P451TAG-R | GTGGTTCCACTCTAAGTGTTTGTTCTGCTCAAGTAATACAGGTACTGG |
| S452TAG-F | GAACAAACACTCCATAGGGAACCACCACGCAGTCAAGGC |
| S452TAG-R | GTGGTGGTTCCCTATGGAGTGTTTGTTCTGCTCAAGTAATACAGGTAC |
| G453TAG-F | CAAACACTCCAAGTTAGACCACCACGCAGTCAAGGCTTCAG |
| G453TAG-R | CTGCGTGGTGGTCTAACTTGGAGTGTTTGTTCTGCTCAAGTAATAGCAGG |
| T454TAG-F | CAAACACTCCAAGTGGATAGACCACGCAGTCAAGGCTTCAGTTTTC |
| T454TAG-R | CTGCGTGGTCTATCCACTTGGAGTGTTTGTTCTGCTCAAGTAATACAGG |
| T455TAG-F | GTGGAACCTAGACGCAGTCAAGGCTTCAGTTTTCTCAG |
| T455TAG-R | GAAGCCTTGACTGCGTCTAGGTTGCACTTGGAGTGTTTGTTCTGCTC |
| T456TAG-F | GTGGAACCACCTAGCAGTCAAGGCTTCAGTTTTCTCAGGCCG |
| T456TAG-R | GAAGCCTTGACTGCTAGGTGGTTCCACTTGGAGTGTTTGTTC |
| Q457TAG-F | GTGGAACCACCACGTAGTCAAGGCTTCAGTTTTCTCAGGCCG |
| Q457TAG-R | GAAGCCTTGACTACGTGGTGGTTCCACTTGGAGTGTTTG |
| S458TAG-F | CACCACGCAGTAGAGGCTTCAGTTTTCTCAGGCCGGAG |
| S458TAG-R | CTGAAGCCTCTACTGCGTGGTGGTTCCACTTGGAGTG |
| R459TAG-F | CACGCAGTCATAGCTTCAGTTTTCTCAGGCCGGAGCG |
| R459TAG-R | GAGAAAACTGAAGCTATGACTGCGTGGTGGTTCCACTTGG |
| VP1-M1L-F | GATTTAAATCAGGTCTCGCTGCCGATGGTTATCTTCCAGATTGGC |
| VP1-M1L-R | CCATCGGCAGCGAGACCTGATTTAAATCATTTATTGTTCAAAGATGCAGTCATCC |
| VP2-T138T-F | GAGGAACCTGTTAAGACCGCTCCGGGAAAAAAGAGGCCGGTAGAG |
| VP2-T138T-R | CCCGGAGCGGTCTTAACAGGTTCCTCAACCAGGCCC |
| VP3-M235L-F | TTCCACATGGCTCGGCGACAGAGTCATCACCACCAG |
| VP3-M235L-R | ACTCTGTCGCCGAGCCATGTGGAATCGCAATGCCAATTTC |

| | |
|---|---|
| VP3-L202L/M203L/211L-R | CTAATACCCTCGCTACAGGCAGTGGCGCACCACTCGCAGACAATAACGAGGGCGCCGACG |
| VP3-L202L/M203L/211L-F | GCGCCACTGCCTGTAGCGAGGGTATTAGTTCCGAGACCAGAGGGGGCTGCTGGTGGCTG |
| RC2-HindIII-F | CGTCAGACGCGGAAGCTTCGATCAAC |
| Cap2-SbfI-R | TTCGATCATTCCTGCAGGTGTAGTTAATGATTAACCCGCC |
| VP1-NheI-F | TTATTTAGCTAGCATGGCTGCCGATGGTTATCTTCCAGATTGGC |
| VP2-ATG-NheI-F | AAGCTGGCTAGCATGGCTCCGGGAAAAAAGAGGCCGG |
| Cap2-AvrII-R | TTATTTACCTAGGGCTGTAGTTAATGATTAACCCGCCATGCTACTTATC |
| SbfI-CMV-F | AGGTACCTTCAACCTGCAGGTTGACATTGATTATTGACTAG |
| Cap2-MluI-R | TAAGAGAATTACGCGTTGTAGTTAATGATTAACCCGCCATGC |

**Production of ncAA containing AAV.**

AAV2 was produced by transfecting HEK293T cells with pHelper, pIDTsmart-*Mb*PylRS-8x*Mm*PyltR-ITR-GFP, and plasmid containing suitable AAV *Rep-Cap* genes in 1:1:1 molar ratio. The plasmids were mixed with polyethyleneimine (Sigma) in serum-free media (DMEM, HyClone) and were incubated at RT for 15 min before adding to HEK293T cells at 70% confluency. AzK (H-L-Lys(EO-N3)-OH, Iris biotech GMBH) was added to the final concentration of 1 mM for the production of AAV2-containing AzK. For a 12 well plate, 1.5 µg total DNA per well was used and AAV2 was harvested 72 h post-transfection using AAVPro Extraction Solution Kit (Takara). In brief, media was removed and cells from each well were harvested and resuspended in 50 µL solution A. After 10 min-incubation, cell debris was pelleted by centrifugation at 12,000 xg for 10 min. The supernatant containing viruses was overlaid in 5 µL solution B. For 15 cm dishes, 57 µg total DNA was used and AAV2 was harvested 120 h post-transfection. Two rounds of freeze-thaw cycles using a dry-ice ethanol bath and a 37 ˚C water bath were used to lyse the cells, followed by centrifugation at 12,000 xg for 5 min to pellet cellular debris. The cell lysate and media were combined and polyethylene glycol (PEG 8000, Fisher Bioreagents) precipitated overnight. AAV2 was pelleted by centrifugation at 5,000 xg for 30 min and resuspended in 2 mL 10% glycerol in PBS. Viruses were titered using the AAVPro titration kit (Takara) and stored at -80˚C for future use, or purified by AVB sepharose (Cytiva) High Performance purification, following the manufacturer's instructions.

**Assaying the infectivity of AAV2.**

Infectivity was assayed as previously described.[1]

**TAMRA labeling, SDS Page, and Western Blotting.**

Approximately $10^{10}$ genome copies of purified virus were incubated 20 µM DBCO-TAMRA (Sigma-Aldrich) at room temperature for 30 min, heated in SDS-loading buffer for 1 min, then analyzed by 10% SDS-PAGE gel. TAMRA fluorescence was imaged with the rhodamine settings on the ChemiDoc MP imaging system (BioRad). Proteins were stained with SYPRO™ Orange

Protein Stain (Thermo Fisher scientific), followed the manufacturer's instructions and imaged with Dylight 540.

AAV production for western blot was done in a 12-well-plate scale experiment. After 72 hours, media was removed, cells from each well were washed once with PBS, and then lysed with 50 µL of CelLytic-M lysis buffer (Sigma; supplemented 0.01% Pierce universal nuclease). After 30 minutes of incubation at RT, 10 µL of cell lysates were heated in SDS-loading buffer and resolved using 10% SDS-PAGE gel. Resolved proteins were transferred from the gel to a PVDF membrane (Life Technologies) using a Trans-Blot Turbo Transfer System (Bio-Rad). The membrane was blocked with blocking solution (5% nonfat milk and 0.1% Tween 20 (Fisher Scientific) in Tris-buffered saline (TBS)) overnight at 4°C. On the next day, blocking solution was removed, and the membrane was incubated with mouse anti-AAV-B1 IgG primary antibody (Progen; 1:4000 dilution) for 2 hours in fresh blocking solution with gentle shaking at RT. The membrane was washed 6 times (10-minute incubation with shaking) with wash solution (0.1% Tween 20 in TBS). Next, the membrane was incubated with chicken anti-mouse IgG secondary antibody-HRP conjugate (Fisher Scientific; 1:3000 dilution) in blocking solution for 1 hour with shaking at RT, and the wash step was repeated. The membrane was developed using SuperSignal West Dura Kit (Fisher Scientific) and incubated for 2 minutes before signal detection by the ChemiDoc MP imaging system (BioRad).

**DBCO-cRGDFC synthesis.** DBCO-cRGDFC was synthesized as previously described.[1] DBCO-PEG24-cRGDFC was synthesized similarly by incubating 9 mM DBCO-PEG24-maleimide (BroadPharm) and 11 mM cRGDFC (Peptides International) in DMF at RT for 2 hours. The reaction completion was verified by HPLC-MS analysis (Agilent Technologies, 1260 Infinity ESI–TOF, Phenomenex, Aeris™ 3.6 µm WIDEPORE XB-C8 column, LC Column 100 x 4.6 mm).

**Re-targeting and flow cytometry.**

Azide-containing virus preparations were labeled with 20 µM DBCO-cRGDFC for varying amounts of time at room temperature and subsequently quenched with 1 mM AzK. $2x10^5$ SK-OV-3 cells were seeded per 12-well plate 24 h prior to infection. When cells had reached desired confluency, virus (2,500 gc/cell) was added to each well along with 5 mM sodium butyrate (Sigma-Aldrich) to enhance the expression of AAV2-encoded transgenes. 48 h post-infection, infectivity was visualized by EGFP expression using a Zeiss Axio Observer fluorescence microscope with an XCite Series 120Q light source and Zeiss filter 44 (excitation 475/40nm, beamsplitter 500nm, emission 530/50nm). To prepare cells for FACS, 200 µL of warm 0.25% trypsin-EDTA solution was added to each well. Plates were incubated at 37 ˚C until cells began to detach from the plate, for about 2 min. 500 µL of ice-cold DMEM + 10% FBS was added to each well to quench the trypsin. Cells were re-suspended by gentle pipetting and transferred to a microcentrifuge tube on ice. An additional 500 µL of DMEM + 10% FBS was used to rinse each well and the rinse was combined with the re-suspended cells. Cells were pelleted by centrifugation at 2,500 xg for five minutes. The supernatant was discarded and cells were gently re-suspended in 500 µL of ice-cold PBS and passed through a 100 µm filter. FACS analysis was performed using a Bio-Rad S3e cell sorter to quantify EGFP fluorescence.

### Plasmid maps

#### pIDTsmart-RC2-WT

Annotation: Rep2, Cap2

cccgtgtaaaacgacggccagtttatctagtcagcttgattctagctgatcgtggaccggaaggtgagccagtgagttgattgcagtccagttacgctggagtctgaggctcgtcctgaatgat
atgcgaccgccggagggttgcgtttgagacgggcgacagatccagtcgcgctgctctcgtcgatccgctagggcggccgctctagaactagtggatcccccggaagatcagaagttccta
ttccgaagttcctattctctagaaagtataggaacttctgatctgcgcagccgccatgccggggttttacgagattgtgattaaggtccccagcgaccttgacgagcatctgcccggcatttctga
cagctttgtgaactgggtggccgagaaggaatgggagttgccgccagattctgacatggatctgaatctgattgagcaggcacccctgaccgtggccgagaagctgcagcgcgactttctg
acggaatggcgccgtgtgagtaaggccccggaggcccttttctttgtgcaatttgagaagggagagagctacttccacatgcacgtgctcgtggaaaccaccggggtgaaatccatggtttt
gggacgtttcctgagtcagattcgcgaaaaactgattcagagaatttaccgcgggatcgagccgactttgccaaactggttcgcggtcacaaagaccagaaatggcgccggaggcggga
acaaggtggtggatgagtgctacatccccaattacttgctccccaaaacccagcctgagctccagtgggcgtggactaatatggaacagtatttaagcgcctgtttgaatctcacggagcgt
aaacggttggtggcgcagcatctgacgcacgtgtcgcagacgcaggagcagaacaaagagaatcagaatcccaattctgatgcgccggtgatcagatcaaaaacttcagccaggtac
atggagctggtcgggtggctcgtggacaaggggattacctcggagaagcagtggatccaggaggaccaggcctcatacatctccttcaatgcggcctccaactcgcggtcccaaatcaa
ggctgccttggacaatgcgggaaagattatgagcctgactaaaaccgcccccgactacctggtgggccagcagcccgtggaggacatttccagcaatcggatttataaaattttggaacta
aacgggtacgatccccaatatgcggcttccgtctttctgggatgggccacgaaaaagttcggcaagaggaacaccatctggctgtttgggcctgcaactaccgggaagaccaacatcgcg
gaggccatagcccacactgtgcccttctacgggtgcgtaaactggaccaatgagaactttcccttcaacgactgtgtcgacaagatggtgatctggtgggaggaggggaagatgaccgcc
aaggtcgtggagtcggccaaagccattctcggaggaagcaaggtgcgcgtggaccagaaatgcaagtcctcggcccagatagacccgactcccgtgatcgtcacctccaacaccaac
atgtgcgccgtgattgacgggaactcaacgaccttcgaacaccagcagccgttgcaagaccggatgttcaaatttgaactcacccgccgtctggatcatgactttgggaaggtcaccaagc
aggaagtcaaagactttttccggtgggcaaaggatcacgtggttgaggtggagcatgaattctacgtcaaaaagggtggagccaagaaaagacccgcccccagtgacgcagatataag
tgagcccaaacgggtgcgcgagtcagttgcgcagccatcgacgtcagacgcggaagcttcgatcaactacgcagacaggtaccaaaacaaatgttctcgtcacgtgggcatgaatctg
atgctgtttccctgcagacaatgcgagagaatgaatcagaattcaaatatctgcttcactcacggacagaaagactgtttagagtgctttcccgtgtcagaatctcaacccgtttctgtcgtcaa
aaaggcgtatcagaaactgtgctacattcatcatatcatgggaaaggtgccagacgcttgcactgcctgcgatctggtcaatgtggatttggatgactgcatctttgaacaataaatgatttaaa
tcaggtatggctgccgatggttatcttccagattggctcgaggacactctctctgaaggaataagacagtggtggaagctcaaacctggcccaccaccaccaaagcccgcagagcggcat
aaggacgacagcaggggtcttgtgcttcctgggtacaagtacctcggacccttcaacggactcgacaagggagagccggtcaacgaggcagacgccgcggccctcgagcacgacaa
agcctacgaccggcagctcgacagcggagacaacccgtacctcaagtacaaccacgccgacgcggagtttcaggagcgccttaaagaagatacgtcttttgggggcaacctcggacg
agcagtcttccaggcgaaaaagagggttcttgaacctctgggcctggttgaggaacctgttaagacggctccgggaaaaaagaggccggtagagcactctcctgtggagccagactcct
cctcgggaaccggaaaggcgggccagcagcctgcaagaaaaagattgaattttggtcagactggagacgcagactcagtacctgacccccagcctctcggacagccaccagcagcc
ccctctggtctgggaactaatacgatggctacaggcagtggcgcaccaatggcagacaataacgagggcgccgacggagtgggtaattcctcgggaaattggcattgcgattccacatg
gatgggcgacagagtcatcaccaccagcacccgaacctgggccctgcccacctacaacaaccacctctacaaacaaatttccagccaatcaggagcctcgaacgacaatcactacttt
ggctacagcaccccttgggggtattttgacttcaacagattccactgccacttttcaccacgtgactggcaaagactcatcaacaacaactggggattccgacccaagagactcaacttcaa
gctctttaacattcaagtcaaagaggtcacgcagaatgacggtacgacgacgattgccaataaccttaccagcacggttcaggtgtttactgactcggagtaccagctcccgtacgtcctcg
gctcggcgcatcaaggatgcctcccgccgttcccagcagacgtcttcatggtgccacagtatggatacctcaccctgaacaacgggagtcaggcagtaggacgctcttcattttactgcctg
gagtactttccttctcagatgctgcgtaccggaaacaactttaccttcagctacacttttgaggacgttcctttccacagcagctacgctcacagccagagtctggaccgtctcatgaatcctctc
atcgaccagtacctgtattacttgagcagaacaaacactccaagtggaaccaccacgcagtcaaggcttcagttttctcaggccggagcgagtgacattcgggaccagtctaggaactgg
cttcctggaccctgttaccgccagcagcgagtatcaaagacatctgcggataacaacaacagtgaatactcgtggactggagctaccaagtaccacctcaatggcagagactctctggtg
aatccgggcccggccatggcaagccacaaggacgatgaagaaaagtttttcctcagagcggggttctcatctttgggaagcaaggctcagagaaaacaaatgtggacattgaaaaggt
catgattacagacgaagaggaaatcaggacaaccaatcccgtggctacggagcagtatggttctgtatctaccaacctccagagaggcaacagacaagcagctaccgcagatgtcaa
cacacaaggcgttcttccaggcatggtctggcaggacagagatgtgtaccttcaggggcccatctgggcaaagattccacacacggacggacattttcacccctctcccctcatgggtgga
ttcggacttaaacaccctcctccacagattctcatcaagaacaccccggtacctgcgaatccttcgaccaccttcagtgcggcaaagtttgcttccttcatcacacagtactccacgggacag
gtcagcgtggagatcgagtgggagctgcagaaggaaaacagcaaacgctggaatcccgaaattcagtacacttccaactacaacaagtctgttaatgtggactttactgtggacactaat
ggcgtgtattcagagcctcgccccattggcaccagatacctgactcgtaatctgtaattgcttgttaatcaataaaccgtttaattcgtttcagttgaactttggtctctgcgtatttctttcttatctagttt
ccatggctacgtagataagtagcatggcgggttaatcattaactacagcccgggcgtttaaacagcgggcggaggggtggagtcgtgacgtgaattacgtcatagggttagggaggtcct
gtattagaggtcacgtgagtgttttgcgacattttgcgacaccatgtggtctcgctgggggggggggcccgagtgagcacgcagggtctccattttgaagcgggaggtttgaacgagcgctg
gcgcgctcactggccgtcgttttacaacgtcgtgactgggaaaaccctggcgttacccaacttaatcgccttgcagcacatccccctttcgccagctggcgtaatagcgaagaggcccgca
ccgatcgcccttcccatgcatcggccgcaaatacctgcaggatccgttttgcgctgcttcgcgatgtacgggccagatatacgcgttgacattgattattgactagttattaatagtaatcaattac
ggggtcattagttcatagcccatatatggagttccgcgttacataacttacggtaaatggcccgcctggctgaccgcccaacgacccccgcccattgacgtcaataatgacgtatgttcccat
agtaacgccaatagggactttccattgacgtcaatgggtggactatttacggtaaactgcccacttggcagtacatcaagtgtatcatatgccaagtacgccccctattgacgtcaatgacggt
aaatggcccgcctggcattatgcccagtacatgaccttatgggactttcctacttggcagtacatctacgtattagtcatcgctattaccatggtgatgcggttttggcagtacatcaatgggcgtg
gatagcggtttgactcacggggatttccaagtctccaccccattgacgtcaatgggagtttgttttggcaccaaaatcaacgggactttccaaaatgtcgtaacaactccgccccattgacgca
aatgggcggtaggcgtgtacggtgggaggtctatataagcagagctctctggctaactagagaacccactgcttactggcttatcgaaattaatacgactcactatagggagacccaagct
ggctaggggtgcgagcggatcgagcagtgtcgatcactactggaccgcgagctgtgctgcgacccgtgatcttacggcattatacgtatgatcggtccacgatcagctagattatctagtca
gcttgatgtcatagctgtttcctgaggctcaatactgaccatttaaatcatacctgacctccatagcagaaagtcaaaagcctccgaccggaggcttttgacttgatcggcacgtaagaggttc
caactttcaccataatgaaataagatcactaccgggcgtatttttttgagttatcgagattttcaggagctaaggaagctaaaatgagccatattcaacgggaaacgtcttgcttgaagccgcga
ttaaattccaacatggatgctgatttatatgggtataaatgggctcgcgataatgtcgggcaatcaggtgcgacaatctatcgattgtatgggaagcccgatgcgccagagttgtttctgaaac
atggcaaaggtagcgttgccaatgatgttacagatgagatggtcaggctaaactggctgacggaatttatgcctcttccgaccatcaagcattttatccgtactcctgatgatgcatggttactc
accactgcgatccaggggaaaacagcattccaggtattagaagaatatcctgattcaggtgaaaatattgttgatgcgctggcagtgttcctgcgccggttgcattcgattcctgtttgtaattgt
ccttttaacggcgatcgcgtatttcgtctcgctcaggcgcaatcacgaatgaataacggtttggttggtgcgagtgattttgatgacgagcgtaatggctggcctgttgaacaagtctggaaaga
aatgcataaactcttgccattctcaccggattcagtcgtcactcatggtgatttctcacttgataaccttatttttgacgaggggaaattaataggttgtattgatgttggacgagtcggaatcgcag

accgataccaggatcttgccatcctatggaactgcctcggtgagttttctccttcattacagaaacggctttttcaaaaatatggtattgataatcctgatatgaataaattgcagtttcacttgatgc
tcgatgagtttttctaatgaggacctaaatgtaatcacctggctcaccttcgggtgggcctttctgcgttgctggcgtttttccataggctccgcccccctgacgagcatcacaaaaatcgatgctc
aagtcagaggtggcgaaacccgacaggactataaagataccaggcgtttccccctggaagctccctcgtgcgctctcctgttccgaccctgccgcttaccggatacctgtccgcctttctccc
ttcgggaagcgtggcgctttctcatagctcacgctgtaggtatctcagttcggtgtaggtcgttcgctccaagctgggctgtgtgcacgaaccccccgttcagcccgaccgctgcgccttatccg
gtaactatcgtcttgagtccaacccggtaagacacgacttatcgccactggcagcagccactggtaacaggattagcagagcgaggtatgtaggcggtgctacagagttcttgaagtggtg
gcctaactacggctacactagaagaacagtatttggtatctgcgctctgctgaagccagttacctcggaaaaagagttggtagctcttgatccggcaaacaaaccaccgctggtagcggtg
gtttttttgtttgcaagcagcagattacgcgcagaaaaaaaggatctcaagaagatcctttgatttctaccgaagaaaggccca

### pIDTsmart-RC2-ΔVP1-CMV-VP1

Annotation: Rep2, Cap2ΔVP1, VP1

cccgtgtaaaacgacggccagtttatctagtcagcttgattctagctgatcgtggaccggaaggtgagccagtgagttgattgcagtccagttacgctggagtctgaggctcgtcctgaatgat
atgcgaccgccggagggttgcgtttgagacgggcgacagatccagtcgcgctgctctcgtcgatccgctagggcggccgctctagaactagtggatcccccggaagatcagaagttccta
ttccgaagttcctattctctagaaagtataggaacttctgatctgcgcagccgccatgccggggttttacgagattgtgattaaggtccccagcgaccttgacgagcatctgcccggcatttctga
cagctttgtgaactgggtggccgagaaggaatgggagttgccgccagattctgacatggatctgaatctgattgagcaggcacccctgaccgtggccgagaagctgcagcgcgactttctg
acggaatggcgccgtgtgagtaaggccccggaggcccttttctttgtgcaatttgagaagggagagagctacttccacatgcacgtgctcgtggaaaccaccggggtgaaatccatggtttt
gggacgtttcctgagtcagattcgcgaaaaactgattcagagaatttaccgcgggatcgagccgactttgccaaactggttcgcggtcacaaagaccagaaatggcgccggaggcggga
acaaggtggtggatgagtgctacatccccaattacttgctccccaaaacccagcctgagctccagtgggcgtggactaatatggaacagtatttaagcgcctgtttgaatctcacggagcgt
aaacggttggtggcgcagcatctgacgcacgtgtcgcagacgcaggagcagaacaaagagaatcagaatcccaattctgatgcgccggtgatcagatcaaaaacttcagccaggtac
atggagctggtcgggtggctcgtggacaaggggattacctcggagaagcagtggatccaggaggaccaggcctcatacatctccttcaatgcggcctccaactcgcggtcccaaatcaa
ggctgccttggacaatgcgggaaagattatgagcctgactaaaaccgcccccgactacctggtgggccagcagcccgtggaggacatttccagcaatcggatttataaaattttggaacta
aacgggtacgatccccaatatgcggcttccgtctttctgggatgggccacgaaaaagttcggcaagaggaacaccatctggctgtttgggcctgcaactaccgggaagaccaacatcgcg
gaggccatagcccacactgtgcccttctacgggtgcgtaaactggaccaatgagaactttcccttcaacgactgtgtcgacaagatggtgatctggtgggaggaggggaagatgaccgcc
aaggtcgtggagtcggccaaagccattctcggaggaagcaaggtgcgcgtggaccagaaatgcaagtcctcggcccagatagacccgactcccgtgatcgtcacctccaacaccaac
atgtgcgccgtgattgacgggaactcaacgaccttcgaacaccagcagccgttgcaagaccggatgttcaaatttgaactcacccgccgtctggatcatgactttgggaaggtcaccaagc
aggaagtcaaagactttttccggtgggcaaaggatcacgtggttgaggtggagcatgaattctacgtcaaaaagggtggagccaagaaaagacccgcccccagtgacgcagatataag
tgagcccaaacgggtgcgcgagtcagttgcgcagccatcgacgtcagacgcggaagcttcgatcaactacgcagacaggtaccaaaacaaatgttctcgtcacgtgggcatgaatctg
atgctgtttccctgcagacaatgcgagagaatgaatcagaattcaaatatctgcttcactcacggacagaaagactgtttagagtgctttcccgtgtcagaatctcaacccgtttctgtcgtcaa
aaaggcgtatcagaaactgtgctacattcatcatatcatgggaaaggtgccagacgcttgcactgcctgcgatctggtcaatgtggatttggatgactgcatctttgaacaataaatgatttaaa
tcaggt**CTC**gctgccgatggttatcttccagattggctcgaggacactctctctgaaggaataagacagtggtggaagctcaaacctggcccaccaccaccaaagcccgcagagcggc
ataaggacgacagcaggggtcttgtgcttcctgggtacaagtacctcggacccttcaacggactcgacaagggagagccggtcaacgaggcagacgccgcggccctcgagcacgac
aaagcctacgaccggcagctcgacagcggagacaacccgtacctcaagtacaaccacgccgacgcggagtttcaggagcgccttaaagaagatacgtcttttgggggcaacctcgga
cgagcagtcttccaggcgaaaaagagggttcttgaacctctgggcctggttgaggaacctgttaagacggctccgggaaaaaagaggccggtagagcactctcctgtggagccagactc
ctcctcgggaaccggaaaggcgggccagcagcctgcaagaaaaagattgaattttggtcagactggagacgcagactcagtacctgacccccagcctctcggacagccaccagcag
ccccctctggtctgggaactaatacgatggctacaggcagtggcgcaccaatggcagacaataacgagggcgccgacggagtgggtaattcctcgggaaattggcattgcgattccaca
tggatgggcgacagagtcatcaccaccagcacccgaacctgggccctgcccacctacaacaaccacctctacaaacaaatttccagccaatcaggagcctcgaacgacaatcactact
ttggctacagcaccccttgggggtattttgacttcaacagattccactgccacttttcaccacgtgactggcaaagactcatcaacaacaactggggattccgacccaagagactcaacttca
agctctttaacattcaagtcaaagaggtcacgcagaatgacggtacgacgacgattgccaataaccttaccagcacggttcaggtgtttactgactcggagtaccagctcccgtacgtcctc
ggctcggcgcatcaaggatgcctcccgccgttcccagcagacgtcttcatggtgccacagtatggatacctcaccctgaacaacgggagtcaggcagtaggacgctcttcattttactgcct
ggagtactttccttctcagatgctgcgtaccggaaacaactttaccttcagctacacttttgaggacgttcctttccacagcagctacgctcacagccagagtctggaccgtctcatgaatcctct
catcgaccagtacctgtattacttgagcagaacaaacactccaagtggaaccaccacgcagtcaaggcttcagttttctcaggccggagcgagtgacattcgggaccagtctaggaactg
gcttcctggaccctgttaccgccagcagcgagtatcaaagacatctgcggataacaacaacagtgaatactcgtggactggagctaccaagtaccacctcaatggcagagactctctggt
gaatccgggcccggccatggcaagccacaaggacgatgaagaaaagtttttttcctcagagcggggttctcatctttgggaagcaaggctcagagaaaacaaatgtggacattgaaaag
gtcatgattacagacgaagaggaaatcaggacaaccaatcccgtggctacggagcagtatggttctgtatctaccaacctccagagaggcaacagacaagcagctaccgcagatgtca
acacacaaggcgttcttccaggcatggtctggcaggacagagatgtgtaccttcaggggcccatctgggcaaagattccacacacggacggacattttcacccctctcccctcatgggtgg
attcggacttaaacaccctcctccacagattctcatcaagaacaccccggtacctgcgaatccttcgaccaccttcagtgcggcaaagtttgcttccttcatcacacagtactccacgggaca
ggtcagcgtggagatcgagtgggagctgcagaaggaaaacagcaaacgctggaatcccgaaattcagtacacttccaactacaacaagtctgttaatgtggactttactgtggacactaa
tggcgtgtattcagagcctcgccccattggcaccagatacctgactcgtaatctgtaattgcttgttaatcaataaaccgtttaattcgtttcagttgaactttggtctctgcgtatttctttcttatctagtt
tccatggctacgtagataagtagcatggcgggttaatcattaactacagcccgggcgtttaaacagcgggcggaggggtggagtcgtgacgtgaattacgtcatagggttagggaggtcct
gtattagaggtcacgtgagtgttttgcgacattttgcgacaccatgtggtctcgctgggggggggggcccgagtgagcacgcagggtctccattttgaagcgggaggtttgaacgagcgctg
gcgcgctcactggccgtcgttttacaacgtcgtgactgggaaaaccctggcgttacccaacttaatcgccttgcagcacatccccctttcgccagctggcgtaatagcgaagaggcccgca
ccgatcgcccttcccatgcatcggccgcaaatacctgcaggatccgttttgcgctgcttcgcgatgtacgggccagatatacgcgttgacattgattattgactagttattaatagtaatcaattac
ggggtcattagttcatagcccatatatggagttccgcgttacataacttacggtaaatggcccgcctggctgaccgcccaacgacccccgcccattgacgtcaataatgacgtatgttcccat
agtaacgccaatagggactttccattgacgtcaatgggtggactatttacggtaaactgcccacttggcagtacatcaagtgtatcatatgccaagtacgccccctattgacgtcaatgacggt
aaatggcccgcctggcattatgcccagtacatgaccttatgggactttcctacttggcagtacatctacgtattagtcatcgctattaccatggtgatgcggttttggcagtacatcaatgggcgtg
gatagcggtttgactcacggggatttccaagtctccaccccattgacgtcaatgggagtttgttttggcaccaaaatcaacgggactttccaaaatgtcgtaacaactccgccccattgacgca
aatgggcggtaggcgtgtacggtgggaggtctatataagcagagctctctggctaactagagaacccactgcttactggcttatcgaaattaatacgactcactatagggagacccaagct
ggctagcatggctgccgatggttatcttccagattggctcgaggacactctctctgaaggaataagacagtggtggaagctcaaacctggcccaccaccaccaaagcccgcagagcggc
ataaggacgacagcaggggtcttgtgcttcctgggtacaagtacctcggacccttcaacggactcgacaagggagagccggtcaacgaggcagacgccgcggccctcgagcacgac
aaagcctacgaccggcagctcgacagcggagacaacccgtacctcaagtacaaccacgccgacgcggagtttcaggagcgccttaaagaagatacgtcttttgggggcaacctcgga

cgagcagtcttccaggcgaaaaagagggttcttgaacctctgggcctggttgaggaacctgttaag**ACC**gctccgggaaaaaagaggccggtagagcactctcctgtggagccagact
cctcctcgggaaccggaaaggcgggccagcagcctgcaagaaaaagattgaattttggtcagactggagacgcagactcagtacctgacccccagcctctcggacagccaccagca
gccccctctggtct**C**ggaactaatac**CCTC**gctacaggcagtggcgcacca**CTC**gcagacaataacgagggcgccgacggagtgggtaattcctcgggaaattggcattgcgattc
cacatgg**CTC**ggcgacagagtcatcaccaccagcacccgaacctgggccctgcccacctacaacaaccacctctacaaacaaatttccagccaatcaggagcctcgaacgacaat
cactactttggctacagcaccccttgggggtattttgacttcaacagattccactgccacttttcaccacgtgactggcaaagactcatcaacaacaactggggattccgacccaagagactc
aacttcaagctctttaacattcaagtcaaagaggtcacgcagaatgacggtacgacgacgattgccaataaccttaccagcacggttcaggtgtttactgactcggagtaccagctcccgta
cgtcctcggctcggcgcatcaaggatgcctcccgccgttcccagcagacgtcttcatggtgccacagtatggatacctcaccctgaacaacgggagtcaggcagtaggacgctcttcatttt
actgcctggagtactttccttctcagatgctgcgtaccggaaacaactttaccttcagctacacttttgaggacgttcctttccacagcagctacgctcacagccagagtctggaccgtctcatga
atcctctcatcgaccagtacctgtattacttgagcagaacaaacactccaagtggaaccaccacgcagtcaaggcttcagttttctcaggccggagcgagtgacattcgggaccagtctag
gaactggcttcctggaccctgttaccgccagcagcgagtatcaaagacatctgcggataacaacaacagtgaatactcgtggactggagctaccaagtaccacctcaatggcagagact
ctctggtgaatccgggcccggccatggcaagccacaaggacgatgaagaaaagtttttcctcagagcggggttctcatctttgggaagcaaggctcagagaaaacaaatgtggacattg
aaaaggtcatgattacagacgaagaggaaatcaggacaaccaatcccgtggctacggagcagtatggttctgtatctaccaacctccagagaggcaacagacaagcagctaccgcag
atgtcaacacacaaggcgttcttccaggcatggtctggcaggacagagatgtgtaccttcaggggcccatctgggcaaagattccacacacggacggacattttcacccctctcccctcatg
ggtggattcggacttaaacaccctcctccacagattctcatcaagaacaccccggtacctgcgaatccttcgaccaccttcagtgcggcaaagtttgcttccttcatcacacagtactccacgg
gacaggtcagcgtggagatcgagtgggagctgcagaaggaaaacagcaaacgctggaatcccgaaattcagtacacttccaactacaacaagtctgttaatgtggactttactgtggac
actaatggcgtgtattcagagcctcgccccattggcaccagatacctgactcgtaatctgtaattgcttgttaatcaataaaccgtttaattcgtttcagttgaactttggtctctgcgtatttctttcttat
ctagtttccatggctacgtagataagtagcatggcgggttaatcattaactacagccctaggggtgcgagcggatcgagcagtgtcgatcactactggaccgcgagctgtgctgcgacccgt
gatcttacggcattatacgtatgatcggtccacgatcagctagattatctagtcagcttgatgtcatagctgtttcctgaggctcaatactgaccatttaaatcatacctgacctccatagcagaaa
gtcaaaagcctccgaccggaggcttttgacttgatcggcacgtaagaggttccaactttcaccataatgaaataagatcactaccgggcgtatttttgagttatcgagattttcaggagctaag
gaagctaaaatgagccatattcaacgggaaacgtcttgcttgaagccgcgattaaattccaacatggatgctgatttatatgggtataaatgggctcgcgataatgtcgggcaatcaggtgc
gacaatctatcgattgtatgggaagcccgatgcgccagagttgtttctgaaacatggcaaaggtagcgttgccaatgatgttacagatgagatggtcaggctaaactggctgacggaatttat
gcctcttccgaccatcaagcattttatccgtactcctgatgatgcatggttactcaccactgcgatccagggaaaacagcattccaggtattagaagaatatcctgattcaggtgaaaatattg
ttgatgcgctggcagtgttcctgcgccggttgcattcgattcctgtttgtaattgtccttttaacggcgatcgcgtatttcgtctcgctcaggcgcaatcacgaatgaataacggtttggttggtgcga
gtgattttgatgacgagcgtaatggctggcctgttgaacaagtctggaaagaaatgcataaactcttgccattctcaccggattcagtcgtcactcatggtgatttctcacttgataaccttatttttg
acgaggggaaattaataggttgtattgatgttggacgagtcggaatcgcagaccgataccaggatcttgccatcctatggaactgcctcggtgagttttctccttcattacagaaacggctttttc
aaaaatatggtattgataatcctgatatgaataaattgcagtttcacttgatgctcgatgagtttttctaatgaggacctaaatgtaatcacctggctcaccttcgggtgggcctttctgcgttgctgg
cgtttttccataggctccgcccccctgacgagcatcacaaaaatcgatgctcaagtcagaggtggcgaaacccgacaggactataaagataccaggcgtttccccctggaagctccctcgt
gcgctctcctgttccgaccctgccgcttaccggatacctgtccgcctttctcccttcgggaagcgtggcgctttctcatagctcacgctgtaggtatctcagttcggtgtaggtcgttcgctccaagc
tgggctgtgtgcacgaaccccccgttcagcccgaccgctgcgccttatccggtaactatcgtcttgagtccaacccggtaagacacgacttatcgccactggcagcagccactggtaacag
gattagcagagcgaggtatgtaggcggtgctacagagttcttgaagtggtggcctaactacggctacactagaagaacagtatttggtatctgcgctctgctgaagccagttacctcggaaa
aagagttggtagctcttgatccggcaaacaaaccaccgctggtagcggtggtttttttgtttgcaagcagcagattacgcgcagaaaaaaaggatctcaagaagatcctttgatttctaccga
agaaaggccca

### pIDTsmart-RC2-ΔVP2-CMV-VP2

Annotation: Rep2, Cap2ΔVP2, VP2

cccgtgtaaaacgacggccagtttatctagtcagcttgattctagctgatcgtggaccggaaggtgagccagtgagttgattgcagtccagttacgctggagtctgaggctcgtcctgaatgat
atgcgaccgccggagggttgcgtttgagacgggcgacagatccagtcgcgctgctctcgtcgatccgctagggcggccgctctagaactagtggatcccccggaagatcagaagttccta
ttccgaagttcctattctctagaaagtataggaacttctgatctgcgcagccgccatgccggggttttacgagattgtgattaaggtccccagcgaccttgacgagcatctgcccggcatttctga
cagctttgtgaactgggtggccgagaaggaatgggagttgccgccagattctgacatggatctgaatctgattgagcaggcacccctgaccgtggccgagaagctgcagcgcgactttctg
acggaatggcgccgtgtgagtaaggccccggaggcccttttctttgtgcaatttgagaagggagagagctacttccacatgcacgtgctcgtggaaaccaccggggtgaaatccatggtttt
gggacgtttcctgagtcagattcgcgaaaaactgattcagagaatttaccgcgggatcgagccgactttgccaaactggttcgcggtcacaaagaccagaaatggcgccggaggcggga
acaaggtggtggatgagtgctacatccccaattacttgctccccaaaacccagcctgagctccagtgggcgtggactaatatggaacagtatttaagcgcctgtttgaatctcacggagcgt
aaacggttggtggcgcagcatctgacgcacgtgtcgcagacgcaggagcagaacaaagagaatcagaatcccaattctgatgcgccggtgatcagatcaaaaacttcagccaggtac
atggagctggtcgggtggctcgtggacaaggggattacctcggagaagcagtggatccaggaggaccaggcctcatacatctccttcaatgcggcctccaactcgcggtcccaaatcaa
ggctgccttggacaatgcgggaaagattatgagcctgactaaaaccgcccccgactacctggtgggccagcagcccgtggaggacatttccagcaatcggatttataaaattttggaacta
aacgggtacgatccccaatatgcggcttccgtctttctgggatgggccacgaaaaagttcggcaagaggaacaccatctggctgtttggcctgcaactaccgggaagaccaacatcgcg
gaggccatagcccacactgtgcccttctacgggtgcgtaaactggaccaatgagaactttcccttcaacgactgtgtcgacaagatggtgatctggtgggaggaggggaagatgaccgcc
aaggtcgtggagtcggccaaagccattctcggaggaagcaaggtgcgcgtggaccagaaatgcaagtcctcggcccagatagacccgactcccgtgatcgtcacctccaacaccaac
atgtgcgccgtgattgacgggaactcaacgaccttcgaacaccagcagccgttgcaagaccggatgttcaaatttgaactcacccgccgtctggatcatgactttgggaaggtcaccaagc
aggaagtcaaagactttttccggtgggcaaaggatcacgtggttgaggtggagcatgaattctacgtcaaaaagggtggagccaagaaaagacccgcccccagtgacgcagatataag
tgagcccaaacgggtgcgcgagtcagttgcgcagccatcgacgtcagacgcggaagcttcgatcaactacgcagacaggtaccaaaacaaatgttctcgtcacgtgggcatgaatctg
atgctgtttccctgcagacaatgcgagagaatgaatcagaattcaaatatctgcttcactcacggacagaaagactgtttagagtgctttcccgtgtcagaatctcaacccgtttctgtcgtcaa
aaaggcgtatcagaaactgtgctacattcatcatatcatgggaaaggtgccagacgcttgcactgcctgcgatctggtcaatgtggatttggatgactgcatctttgaacaataaatgatttaaa
tcaggtatggctgccgatggttatcttccagattggctcgaggacactctctctgaaggaataagacagtggtggaagctcaaacctggcccaccaccaccaaagcccgcagagcggcat
aaggacgacagcaggggtcttgtgcttcctgggtacaagtacctcggacccttcaacggactcgacaagggagagccggtcaacgaggcagacgccgcggccctcgagcacgacaa
agcctacgaccggcagctcgacagcggagacaacccgtacctcaagtacaaccacgccgacgcggagtttcaggagcgccttaaagaagatacgtcttttgggggcaacctcggacg
agcagtcttccaggcgaaaaagagggttcttgaacctctgggcctggttgaggaacctgttaag**ACC**gctccgggaaaaaagaggccggtagagcactctcctgtggagccagactcc

tcctcgggaaccggaaaggcgggccagcagcctgcaagaaaaagattgaattttggtcagactggagacgcagactcagtacctgacccccagcctctcggacagccaccagcagc
cccctctggtctgggaactaatacgatggctacaggcagtggcgcaccaatggcagacaataacgagggcgccgacggagtgggtaattcctcgggaaattggcattgcgattccacat
ggatgggcgacagagtcatcaccaccagcacccgaacctgggccctgcccacctacaacaaccacctctacaaacaaatttccagccaatcaggagcctcgaacgacaatcactactt
tggctacagcaccccttgggggtattttgacttcaacagattccactgccacttttcaccacgtgactggcaaagactcatcaacaacaactggggattccgacccaagagactcaacttca
agctctttaacattcaagtcaaagaggtcacgcagaatgacggtacgacgacgattgccaataaccttaccagcacggttcaggtgtttactgactcggagtaccagctcccgtacgtcctc
ggctcggcgcatcaaggatgcctcccgccgttcccagcagacgtcttcatggtgccacagtatggatacctcaccctgaacaacgggagtcaggcagtaggacgctcttcattttactgcct
ggagtactttccttctcagatgctgcgtaccggaaacaactttaccttcagctacacttttgaggacgttcctttccacagcagctacgctcacagccagagtctggaccgtctcatgaatcctct
catcgaccagtacctgtattacttgagcagaacaaacactccaagtggaaccaccacgcagtcaaggcttcagttttctcaggccggagcgagtgacattcgggaccagtctaggaactg
gcttcctggaccctgttaccgccagcagcgagtatcaaagacatctgcggataacaacaacagtgaatactcgtggactggagctaccaagtaccacctcaatggcagagactctctggt
gaatccgggcccggccatggcaagccacaaggacgatgaagaaaagtttttcctcagagcggggttctcatctttgggaagcaaggctcagagaaaacaaatgtggacattgaaaag
gtcatgattacagacgaagaggaaatcaggacaaccaatcccgtggctacggagcagtatggttctgtatctaccaacctccagagaggcaacagacaagcagctaccgcagatgtca
acacacaaggcgttcttccaggcatggtctggcaggacagagatgtgtaccttcaggggcccatctgggcaaagattccacacacggacggacattttcacccctctcccctcatgggtgg
attcggacttaaacaccctcctccacagattctcatcaagaacaccccggtacctgcgaatccttcgaccaccttcagtgcggcaaagtttgcttccttcatcacacagtactccacgggaca
ggtcagcgtggagatcgagtgggagctgcagaaggaaaacagcaaacgctggaatcccgaaattcagtacacttccaactacaacaagtctgttaatgtggactttactgtggacactaa
tggcgtgtattcagagcctcgccccattggcaccagatacctgactcgtaatctgtaattgcttgttaatcaataaaccgtttaattcgtttcagttgaactttggtctctgcgtatttctttcttatctagtt
tccatggctacgtagataagtagcatggcgggttaatcattaactacagcccgggcgtttaaacagcgggcggaggggtggagtcgtgacgtgaattacgtcatagggttagggaggtcct
gtattagaggtcacgtgagtgttttgcgacattttgcgacaccatgtggtctcgctgggggggggggcccgagtgagcacgcagggtctccattttgaagcgggaggtttgaacgagcgctg
gcgcgctcactggccgtcgttttacaacgtcgtgactgggaaaaccctggcgttacccaacttaatcgccttgcagcacatccccctttcgccagctggcgtaatagcgaagaggcccgca
ccgatcgcccttcccatgcatcggccgcaaatacctgcaggatccgttttgcgctgcttcgcgatgtacgggccagatatacgcgttgacattgattattgactagttattaatagtaatcaattac
ggggtcattagttcatagcccatatatggagttccgcgttacataacttacggtaaatggcccgcctggctgaccgcccaacgacccccgcccattgacgtcaataatgacgtatgttcccat
agtaacgccaatagggactttccattgacgtcaatgggtggactatttacggtaaactgcccacttggcagtacatcaagtgtatcatatgccaagtacgccccctattgacgtcaatgacggt
aaatggcccgcctggcattatgcccagtacatgaccttatgggactttcctacttggcagtacatctacgtattagtcatcgctattaccatggtgatgcggttttggcagtacatcaatgggcgtg
gatagcggtttgactcacggggatttccaagtctccaccccattgacgtcaatgggagtttgttttggcaccaaaatcaacgggactttccaaaatgtcgtaacaactccgccccattgacgca
aatgggcggtaggcgtgtacggtgggaggtctatataagcagagctctctggctaactagagaacccactgcttactggcttatcgaaattaatacgactcactatagggagacccaagct
ggctagc**ATG**gctccgggaaaaaagaggccggtagagcactctcctgtggagccagactcctcctcgggaaccggaaaggcgggccagcagcctgcaagaaaaagattgaatttt
ggtcagactggagacgcagactcagtacctgacccccagcctctcggacagccaccagcagccccctctggtct**C**ggaactaatac**CCTC**gctacaggcagtggcgcacca**CTC**
gcagacaataacgagggcgccgacggagtgggtaattcctcgggaaattggcattgcgattccacatgg**CTC**ggcgacagagtcatcaccaccagcacccgaacctgggccctgcc
cacctacaacaaccacctctacaaacaaatttccagccaatcaggagcctcgaacgacaatcactactttggctacagcaccccttgggggtattttgacttcaacagattccactgccactt
ttcaccacgtgactggcaaagactcatcaacaacaactggggattccgacccaagagactcaacttcaagctctttaacattcaagtcaaagaggtcacgcagaatgacggtacgacga
cgattgccaataaccttaccagcacggttcaggtgtttactgactcggagtaccagctcccgtacgtcctcggctcggcgcatcaaggatgcctcccgccgttcccagcagacgtcttcatgg
tgccacagtatggatacctcaccctgaacaacgggagtcaggcagtaggacgctcttcattttactgcctggagtactttccttctcagatgctgcgtaccggaaacaactttaccttcagctac
acttttgaggacgttcctttccacagcagctacgctcacagccagagtctggaccgtctcatgaatcctctcatcgaccagtacctgtattacttgagcagaacaaacactccaagtggaacc
accacgcagtcaaggcttcagttttctcaggccggagcgagtgacattcgggaccagtctaggaactggcttcctggaccctgttaccgccagcagcgagtatcaaagacatctgcggata
acaacaacagtgaatactcgtggactggagctaccaagtaccacctcaatggcagagactctctggtgaatccgggcccggccatggcaagccacaaggacgatgaagaaaagtttttt
cctcagagcggggttctcatctttgggaagcaaggctcagagaaaacaaatgtggacattgaaaaggtcatgattacagacgaagaggaaatcaggacaaccaatcccgtggctacgg
agcagtatggttctgtatctaccaacctccagagaggcaacagacaagcagctaccgcagatgtcaacacacaaggcgttcttccaggcatggtctggcaggacagagatgtgtaccttc
aggggcccatctgggcaaagattccacacacggacggacattttcacccctctcccctcatgggtggattcggacttaaacaccctcctccacagattctcatcaagaacaccccggtacct
gcgaatccttcgaccaccttcagtgcggcaaagtttgcttccttcatcacacagtactccacgggacaggtcagcgtggagatcgagtgggagctgcagaaggaaaacagcaaacgctg
gaatcccgaaattcagtacacttccaactacaacaagtctgttaatgtggactttactgtggacactaatggcgtgtattcagagcctcgccccattggcaccagatacctgactcgtaatctgt
aattgcttgttaatcaataaaccgtttaattcgtttcagttgaactttggtctctgcgtatttctttcttatctagtttccatggctacgtagataagtagcatggcgggttaatcattaactacagccctag
gggtgcgagcggatcgagcagtgtcgatcactactggaccgcgagctgtgctgcgacccgtgatcttacggcattatacgtatgatcggtccacgatcagctagattatctagtcagcttgat
gtcatagctgtttcctgaggctcaatactgaccatttaaatcatacctgacctccatagcagaaagtcaaaagcctccgaccggaggcttttgacttgatcggcacgtaagaggttccaactttc
accataatgaaataagatcactaccgggcgtatttttgagttatcgagattttcaggagctaaggaagctaaaatgagccatattcaacgggaaacgtcttgcttgaagccgcgattaaattc
caacatggatgctgatttatatgggtataaatgggctcgcgataatgtcgggcaatcaggtgcgacaatctatcgattgtatgggaagcccgatgcgccagagttgtttctgaaacatggcaa
aggtagcgttgccaatgatgttacagatgagatggtcaggctaaactggctgacggaatttatgcctcttccgaccatcaagcattttatccgtactcctgatgatgcatggttactcaccactgc
gatcccagggaaaacagcattccaggtattagaagaatatcctgattcaggtgaaaatattgttgatgcgctggcagtgttcctgcgccggttgcattcgattcctgtttgtaattgtccttttaacg
gcgatcgcgtatttcgtctcgctcaggcgcaatcacgaatgaataacggtttggttggtgcgagtgattttgatgacgagcgtaatggctggcctgttgaacaagtctggaaagaaatgcata
aactcttgccattctcaccggattcagtcgtcactcatggtgatttctcacttgataaccttatttttgacgaggggaaattaataggttgtattgatgttggacgagtcggaatcgcagaccgatac
caggatcttgccatcctatggaactgcctcggtgagttttctccttcattacagaaacggctttttcaaaaatatggtattgataatcctgatatgaataaattgcagtttcacttgatgctcgatgagt
tttctaatgaggacctaaatgtaatcacctggctcaccttcgggtgggcctttctgcgttgctggcgtttttccataggctccgcccccctgacgagcatcacaaaaatcgatgctcaagtcaga
ggtggcgaaacccgacaggactataaagataccaggcgtttccccctggaagctccctcgtgcgctctcctgttccgaccctgccgcttaccggatacctgtccgcctttctcccttcgggaa
gcgtggcgctttctcatagctcacgctgtaggtatctcagttcggtgtaggtcgttcgctccaagctgggctgtgtgcacgaaccccccgttcagcccgaccgctgcgccttatccggtaactat
cgtcttgagtccaacccggtaagacacgacttatcgccactggcagcagccactggtaacaggattagcagagcgaggtatgtaggcggtgctacagagttcttgaagtggtggcctaact
acggctacactagaagaacagtatttggtatctgcgctctgctgaagccagttacctcggaaaaagagttggtagctcttgatccggcaaacaaaccaccgctggtagcggtggtttttttgttt
gcaagcagcagattacgcgcagaaaaaaaggatctcaagaagatcctttgatttttctaccgaagaaaggccca

**pIDTsmart-RC2-ΔVP1,2-CMV-VP1,2**

Annotation: Rep2, Cap2ΔVP1,2, VP2, VP1

cccgtgtaaaacgacggccagtttatctagtcagcttgattctagctgatcgtggaccggaaggtgagccagtgagttgattgcagtccagttacgctggagtctgaggctcgtcctgaatgat
atgcgaccgccggagggttgcgtttgagacgggcgacagatccagtcgcgctgctctcgtcgatccgctagggcggccgctctagaactagtggatcccccggaagatcagaagttccta
ttccgaagttcctattctctagaaagtataggaacttctgatctgcgcagccgccatgccggggttttacgagattgtgattaaggtccccagcgaccttgacgagcatctgcccggcatttctga
cagctttgtgaactgggtggccgagaaggaatgggagttgccgccagattctgacatggatctgaatctgattgagcaggcacccctgaccgtggccgagaagctgcagcgcgactttctg
acggaatggcgccgtgtgagtaaggccccggaggcccttttctttgtgcaatttgagaagggagagagctacttccacatgcacgtgctcgtggaaaccaccggggtgaaatccatggtttt
gggacgtttcctgagtcagattcgcgaaaaactgattcagagaatttaccgcgggatcgagccgactttgccaaactggttcgcggtcacaaagaccagaaatggcgccggaggcggga
acaaggtggtggatgagtgctacatccccaattacttgctccccaaaacccagcctgagctccagtgggcgtggactaatatggaacagtatttaagcgcctgtttgaatctcacggagcgt
aaacggttggtggcgcagcatctgacgcacgtgtcgcagacgcaggagcagaacaaagagaatcagaatcccaattctgatgcgccggtgatcagatcaaaaacttcagccaggtac
atggagctggtcgggtggctcgtggacaaggggattacctcggagaagcagtggatccaggaggaccaggcctcatacatctccttcaatgcggcctccaactcgcggtcccaaatcaa
ggctgccttggacaatgcgggaaagattatgagcctgactaaaaccgcccccgactacctggtgggccagcagcccgtggaggacatttccagcaatcggatttataaaattttggaacta
aacgggtacgatccccaatatgcggcttccgtctttctgggatgggccacgaaaaagttcggcaagaggaacaccatctggctgtttggcctgcaactaccgggaagaccaacatcgcg
gaggccatagcccacactgtgcccttctacgggtgcgtaaactggaccaatgagaactttcccttcaacgactgtgtcgacaagatggtgatctggtgggaggaggggaagatgaccgcc
aaggtcgtggagtcggccaaagccattctcggaggaagcaaggtgcgcgtggaccagaaatgcaagtcctcggcccagatagacccgactcccgtgatcgtcacctccaacaccaac
atgtgcgccgtgattgacgggaactcaacgaccttcgaacaccagcagccgttgcaagaccggatgttcaaatttgaactcacccgccgtctggatcatgactttgggaaggtcaccaagc
aggaagtcaaagactttttccggtgggcaaaggatcacgtggttgaggtggagcatgaattctacgtcaaaaagggtggagccaagaaaagacccgcccccagtgacgcagatataag
tgagcccaaacgggtgcgcgagtcagttgcgcagccatcgacgtcagacgcggaagcttcgatcaactacgcagacaggtaccaaaacaaatgttctcgtcacgtgggcatgaatctg
atgctgtttccctgcagacaatgcgagagaatgaatcagaattcaaatatctgcttcactcacggacagaaagactgtttagagtgctttcccgtgtcagaatctcaacccgtttctgtcgtcaa
aaaggcgtatcagaaactgtgctacattcatcatatcatgggaaaggtgccagacgcttgcactgcctgcgatctggtcaatgtggatttggatgactgcatctttgaacaataaatgatttaaa
tcaggt**CTC**gctgccgatggttatcttccagattggctcgaggacactctctctgaaggaataagacagtggtggaagctcaaacctggcccaccaccaccaaagcccgcagagcggc
ataaggacgacagcaggggtcttgtgcttcctgggtacaagtacctcggacccttcaacggactcgacaagggagagccggtcaacgaggcagacgccgcggccctcgagcacgac
aaagcctacgaccggcagctcgacagcggagacaacccgtacctcaagtacaaccacgccgacgcggagtttcaggagcgccttaaagaagatacgtcttttgggggcaacctcgga
cgagcagtcttccaggcgaaaaagagggttcttgaacctctgggcctggttgaggaacctgttaag**ACC**gctccgggaaaaaagaggccggtagagcactctcctgtggagccagact
cctcctcgggaaccggaaaggcgggccagcagcctgcaagaaaaagattgaattttggtcagactggagacgcagactcagtacctgacccccagcctctcggacagccaccagca
gcccccctctggtctgggaactaatacgatggctacaggcagtggcgcaccaatggcagacaataacgagggcgccgacggagtgggtaattcctcgggaaattggcattgcgattccac
atggatgggcgacagagtcatcaccaccagcacccgaacctgggccctgcccacctacaacaaccacctctacaaacaaatttccagccaatcaggagcctcgaacgacaatcacta
ctttggctacagcaccccttgggggtattttgacttcaacagattccactgccacttttcaccacgtgactggcaaagactcatcaacaacaactggggattccgacccaagagactcaacttc
aagctctttaacattcaagtcaaagaggtcacgcagaatgacggtacgacgacgattgccaataaccttaccagcacggttcaggtgtttactgactcggagtaccagctcccgtacgtcct
cggctcggcgcatcaaggatgcctcccgccgttcccagcagacgtcttcatggtgccacagtatggatacctcaccctgaacaacgggagtcaggcagtaggacgctcttcattttactgcc
tggagtactttccttctcagatgctgcgtaccggaaacaactttaccttcagctacacttttgaggacgttcctttccacagcagctacgctcacagccagagtctggaccgtctcatgaatcctct
catcgaccagtacctgtattacttgagcagaacaaacactccaagtggaaccaccacgcagtcaaggcttcagttttctcaggccggagcgagtgacattcgggaccagtctaggaactg
gcttcctggaccctgttaccgccagcagcgagtatcaaagacatctgcggataacaacaacagtgaatactcgtggactggagctaccaagtaccacctcaatggcagagactctctggt
gaatccgggcccggccatggcaagccacaaggacgatgaagaaaagtttttcctcagagcggggttctcatctttgggaagcaaggctcagagaaaacaaatgtggacattgaaaag
gtcatgattacagacgaagaggaaatcaggacaaccaatcccgtggctacggagcagtatggttctgtatctaccaacctccagagaggcaacagacaagcagctaccgcagatgtca
acacacaaggcgttcttccaggcatggtctggcaggacagagatgtgtaccttcaggggcccatctgggcaaagattccacacacggacggacattttcacccctctcccctcatgggtgg
attcggacttaaacaccctcctccacagattctcatcaagaacaccccggtacctgcgaatccttcgaccaccttcagtgcggcaaagtttgcttccttcatcacacagtactccacgggaca
ggtcagcgtggagatcgagtgggagctgcagaaggaaaacagcaaacgctggaatcccgaaattcagtacacttccaactacaacaagtctgttaatgtggactttactgtggacactaa
tggcgtgtattcagagcctcgccccattggcaccagatacctgactcgtaatctgtaattgcttgttaatcaataaaccgtttaattcgtttcagttgaactttggtctctgcgtatttctttcttatctagtt
tccatggctacgtagataagtagcatggcgggttaatcattaactacacctgcaggttgacattgattattgactagttattaatagtaatcaattacggggtcattagttcatagcccatatatgg
agttccgcgttacataacttacggtaaatggcccgcctggctgaccgcccaacgacccccgcccattgacgtcaataatgacgtatgttcccatagtaacgccaatagggactttccattgac
gtcaatgggtggactatttacggtaaactgcccacttggcagtacatcaagtgtatcatatgccaagtacgccccctattgacgtcaatgacggtaaatggcccgcctggcattatgcccagt
acatgaccttatgggactttcctacttggcagtacatctacgtattagtcatcgctattaccatggtgatgcggttttggcagtacatcaatgggcgtggatagcggtttgactcacggggatttcc
aagtctccaccccattgacgtcaatgggagtttgttttggcaccaaaatcaacgggactttccaaaatgtcgtaacaactccgccccattgacgcaaatgggcggtaggcgtgtacggtggg
aggtctatataagcagagctctctggctaactagagaacccactgcttactggcttatcgaaattaatacgactcactatagggagacccaagctggctagc**atg**gctccgggaaaaaag
aggccggtagagcactctcctgtggagccagactcctcctcgggaaccggaaaggcgggccagcagcctgcaagaaaaagattgaattttggtcagactggagacgcagactcagta
cctgacccccagcctctcggacagccaccagcagcccccctctggtct**C**ggaactaatac**CCTC**gctacaggcagtggcgcacca**CTC**gcagacaataacgagggcgccgacgg
agtgggtaattcctcgggaaattggcattgcgattccacatgg**CTC**ggcgacagagtcatcaccaccagcacccgaacctgggccctgcccacctacaacaaccacctctacaaaca
aatttccagccaatcaggagcctcgaacgacaatcactactttggctacagcaccccttgggggtattttgacttcaacagattccactgccacttttcaccacgtgactggcaaagactcatc
aacaacaactggggattccgacccaagagactcaacttcaagctctttaacattcaagtcaaagaggtcacgcagaatgacggtacgacgacgattgccaataaccttaccagcacggt
tcaggtgtttactgactcggagtaccagctcccgtacgtcctcggctcggcgcatcaaggatgcctcccgccgttcccagcagacgtcttcatggtgccacagtatggatacctcaccctgaa
caacgggagtcaggcagtaggacgctcttcattttactgcctggagtactttccttctcagatgctgcgtaccggaaacaactttaccttcagctacacttttgaggacgttcctttccacagcag
ctacgctcacagccagagtctggaccgtctcatgaatcctctcatcgaccagtacctgtattacttgagcagaacaaacactccaagtggaaccaccacgcagtcaaggcttcagttttctca
ggccggagcgagtgacattcgggaccagtctaggaactggcttcctggaccctgttaccgccagcagcgagtatcaaagacatctgcggataacaacaacagtgaatactcgtggactg
gagctaccaagtaccacctcaatggcagagactctctggtgaatccgggcccggccatggcaagccacaaggacgatgaagaaaagtttttcctcagagcggggttctcatctttggga
agcaaggctcagagaaaacaaatgtggacattgaaaaggtcatgattacagacgaagaggaaatcaggacaaccaatcccgtggctacggagcagtatggttctgtatctaccaacct
ccagagaggcaacagacaagcagctaccgcagatgtcaacacacaaggcgttcttccaggcatggtctggcaggacagagatgtgtaccttcaggggcccatctgggcaaagattcc
acacacggacggacattttcacccctctcccctcatgggtggattcggacttaaacaccctcctccacagattctcatcaagaacaccccggtacctgcgaatccttcgaccaccttcagtgc
ggcaaagtttgcttccttcatcacacagtactccacgggacaggtcagcgtggagatcgagtgggagctgcagaaggaaaacagcaaacgctggaatcccgaaattcagtacacttcca
actacaacaagtctgttaatgtggactttactgtggacactaatggcgtgtattcagagcctcgccccattggcaccagatacctgactcgtaatctgtaattgcttgttaatcaataaaccgttta
attcgtttcagttgaactttggtctctgcgtatttctttcttatctagtttccatggctacgtagataagtagcatggcgggttaatcattaactacaacgcgttgacattgattattgactagttattaata
gtaatcaattacggggtcattagttcatagcccatatatggagttccgcgttacataacttacggtaaatggcccgcctggctgaccgcccaacgacccccgcccattgacgtcaataatgac
gtatgttcccatagtaacgccaatagggactttccattgacgtcaatgggtggactatttacggtaaactgcccacttggcagtacatcaagtgtatcatatgccaagtacgccccctattgacg

tcaatgacggtaaatggcccgcctggcattatgcccagtacatgaccttatgggactttcctacttggcagtacatctacgtattagtcatcgctattaccatggtgatgcggttttggcagtacat
caatgggcgtggatagcggtttgactcacggggatttccaagtctccaccccattgacgtcaatgggagtttgttttggcaccaaaatcaacgggactttccaaaatgtcgtaacaactccgc
cccattgacgcaaatgggcggtaggcgtgtacggtgggaggtctatataagcagagctctctggctaactagagaacccactgcttactggcttatcgaaattaatacgactcactataggg
agacccaagctggctagcatggctgccgatggttatcttccagattggctcgaggacactctctctgaaggaataagacagtggtggaagctcaaacctggcccaccaccaccaaagcc
cgcagagcggcataaggacgacagcaggggtcttgtgcttcctgggtacaagtacctcggacccttcaacggactcgacaagggagagccggtcaacgaggcagacgccgcggccc
tcgagcacgacaaagcctacgaccggcagctcgacagcggagacaacccgtacctcaagtacaaccacgccgacgcggagtttcaggagcgccttaaagaagatacgtcttttgggg
gcaacctcggacgagcagtcttccaggcgaaaaagagggttcttgaacctctgggcctggttgaggaacctgttaag**ACC**gctccgggaaaaaagaggccggtagagcactctcctgt
ggagccagactcctcctcgggaaccggaaaggcgggccagcagcctgcaagaaaaagattgaattttggtcagactggagacgcagactcagtacctgacccccagcctctcggaca
gccaccagcagccccctctggtct**C**ggaactaatac**CCTC**gctacaggcagtggcgcacca**CTC**gcagacaataacgagggcgccgacggagtgggtaattcctcgggaaattg
gcattgcgattccacatgg**CTC**ggcgacagagtcatcaccaccagcacccgaacctgggccctgcccacctacaacaaccacctctacaaacaaatttccagccaatcaggagcctc
gaacgacaatcactactttggctacagcaccccttgggggtattttgacttcaacagattccactgccacttttcaccacgtgactggcaaagactcatcaacaacaactggggattccgacc
caagagactcaacttcaagctctttaacattcaagtcaaagaggtcacgcagaatgacggtacgacgacgattgccaataaccttaccagcacggttcaggtgtttactgactcggagtac
cagctcccgtacgtcctcggctcggcgcatcaaggatgcctcccgccgttcccagcagacgtcttcatggtgccacagtatggatacctcaccctgaacaacgggagtcaggcagtagga
cgctcttcattttactgcctggagtactttccttctcagatgctgcgtaccggaaacaactttaccttcagctacacttttgaggacgttcctttccacagcagctacgctcacagccagagtctgga
ccgtctcatgaatcctctcatcgaccagtacctgtattacttgagcagaacaaacactccaagtggaaccaccacgcagtcaaggcttcagttttctcaggccggagcgagtgacattcggg
accagtctaggaactggcttcctggaccctgttaccgccagcagcgagtatcaaagacatctgcggataacaacaacagtgaatactcgtggactggagctaccaagtaccacctcaatg
gcagagactctctggtgaatccgggcccggccatggcaagccacaaggacgatgaagaaaagtttttcctcagagcggggttctcatctttgggaagcaaggctcagagaaaacaaat
gtggacattgaaaaggtcatgattacagacgaagaggaaatcaggacaaccaatcccgtggctacggagcagtatggttctgtatctaccaacctccagagaggcaacagacaagca
gctaccgcagatgtcaacacacaaggcgttcttccaggcatggtctggcaggacagagatgtgtaccttcaggggcccatctgggcaaagattccacacacggacggacattttcacccc
tctcccctcatgggtggattcggacttaaacaccctcctccacagattctcatcaagaacaccccggtacctgcgaatccttcgaccaccttcagtgcggcaaagtttgcttccttcatcacaca
gtactccacgggacaggtcagcgtggagatcgagtgggagctgcagaaggaaaacagcaaacgctggaatcccgaaattcagtacacttccaactacaacaagtctgttaatgtggac
tttactgtggacactaatggcgtgtattcagagcctcgccccattggcaccagatacctgactcgtaatctgtaattgcttgttaatcaataaaccgtttaattcgtttcagttgaactttggtctctgc
gtatttctttcttatctagtttccatggctacgtagataagtagcatggcgggttaatcattaactacagccctaggggtgcgagcggatcgagcagtgtcgatcactactggaccgcgagctgt
gctgcgacccgtgatcttacggcattatacgtatgatcggtccacgatcagctagattatctagtcagcttgatgtcatagctgtttcctgaggctcaatactgaccatttaaatcatacctgacctc
catagcagaaagtcaaaagcctccgaccggaggcttttgacttgatcggcacgtaagaggttccaactttcaccataatgaaataagatcactaccgggcgtatttttttgagttatcgagatttt
caggagctaaggaagctaaaatgagccatattcaacgggaaacgtcttgcttgaagccgcgattaaattccaacatggatgctgatttatatgggtataaatgggctcgcgataatgtcggg
caatcaggtgcgacaatctatcgattgtatgggaagcccgatgcgccagagttgtttctgaaacatggcaaaggtagcgttgccaatgatgttacagatgagatggtcaggctaaactggct
gacggaatttatgcctcttccgaccatcaagcattttatccgtactcctgatgatgcatggttactcaccactgcgatccccagggaaaacagcattccaggtattagaagaatatcctgattcag
gtgaaaatattgttgatgcgctggcagtgttcctgcgccggttgcattcgattcctgtttgtaattgtccttttaacggcgatcgcgtatttcgtctcgctcaggcgcaatcacgaatgaataacggttt
ggttggtgcgagtgattttgatgacgagcgtaatggctggcctgttgaacaagtctggaaagaaatgcataaactcttgccattctcaccggattcagtcgtcactcatggtgatttctcacttgat
aaccttatttttgacgaggggaaattaataggttgtattgatgttggacgagtcggaatcgcagaccgataccaggatcttgccatcctatggaactgcctcggtgagttttctccttcattacaga
aacggctttttcaaaaatatggtattgataatcctgatatgaataaattgcagtttcacttgatgctcgatgagtttttctaatgaggacctaaatgtaatcacctggctcaccttcgggtgggcctttc
tgcgttgctggcgtttttccataggctccgcccccctgacgagcatcacaaaaatcgatgctcaagtcagaggtggcgaaacccgacaggactataaagataccaggcgtttccccctgga
agctccctcgtgcgctctcctgttccgaccctgccgcttaccggatacctgtccgcctttctcccttcgggaagcgtggcgctttctcatagctcacgctgtaggtatctcagttcggtgtaggtcgtt
cgctccaagctgggctgtgtgcacgaaccccccgttcagcccgaccgctgcgccttatccggtaactatcgtcttgagtccaacccggtaagacacgacttatcgccactggcagcagcca
ctggtaacaggattagcagagcgaggtatgtaggcggtgctacagagttcttgaagtggtggcctaactacggctacactagaagaacagtatttggtatctgcgctctgctgaagccagtta
cctcggaaaaagagttggtagctcttgatccggcaaacaaaccaccgctggtagcggtggtttttttgtttgcaagcagcagattacgcgcagaaaaaaaggatctcaagaagatcctttga
ttttctaccgaagaaaggccca

### pIDTsmart-MbPylRS-8xPyltR-ITR-GFP

Annotation: AAV2 ITR, GFP, MmPyltR, MbPylRS

cccgtgtaaaacgacggccagtttatctagtcagcttgattctagctgatcgtggaccggaaggtgagccagtgagttgattgcagtccagttacgctggagtctgaggctcgtcctgaatgat
atgcgaccgccggagggttgcgtttgagacgggcgacagatccagtcgcgctgctctcgtcgatccgctagggcggccgcaaatacctgcaggcagctgcgcgctcgctcgctcactga
ggccgcccgggcaaagcccgggcgtcgggcgacctttggtcgcccggcctcagtgagcgagcgagcgcgcagagagggagtggccaactccatcactaggggttcctgcggccgca
cgcgtggagctagttattaatagtaatcaattacggggtcattagttcatagcccatatatggagttccgcgttacataacttacggtaaatggcccgcctggctgaccgcccaacgacccccg
cccattgacgtcaataatgacgtatgttcccatagtaacgtcaatagggactttccattgacgtcaatgggtggagtatttacggtaaactgcccacttggcagtacatcaagtgtatcatatgcc
aagtacgccccctattgacgtcaatgacggtaaatggcccgcctggcattatgcccagtacatgaccttatgggactttcctacttggcagtacatctacgtattagtcatcgctattaccatggt
gatgcggttttggcagtacatcaatgggcgtggatagcggtttgactcacggggatttccaagtctccaccccattgacgtcaatgggagtttgttttgcaccaaaatcaacgggactttccaaa
atgtcgtaacaactccgccccattgacgcaaatgggcggtaggcgtgtacggtgggaggtctatataagcagagctcgtttagtgaaccgtcagatcgcctggagacgccatccacgctgt
tttgacctccatagaagacaccgggaccgatccagcctccgcggattcgaatcccggccgggaacggtgcattggaacgcggattccccgtgccaagagtgacgtaagtaccgcctata
gagtctataggcccacaaaaaatgctttcttcttttaatatactttttttgtttatcttatttctaatactttccctaatctctttctttcagggcaataatgatacaatgtatcatgcctctttgcaccattctaaa
gaataacagtgataatttctgggttaaggcaatagcaatatttctgcatataaatatttctgcatataaattgtaactgatgtaagaggtttcatattgctaatagcagctacaatccagctaccatt
ctgcttttattttatggttgggataaggctggattattctgagtccaagctaggcccttttgctaatcatgttcatacctcttatcttcctcccacagctcctgggcaacgtgctggtctgtgtgctggccc
atcactttggcaaagaattgggattcgaacatcgattgaattctgaatggtgagcaagggcgaggagctgttcaccggggtggtgcccatcctggtcgagctggacggcgacgtaaacgg
ccacaagttcagcgtgtccggcgagggcgagggcgatgccacctacggcaagctgaccctgaagttcatctgcaccaccggcaagctgcccgtgccctggcccaccctcgtgaccacc
ctgacctacggcgtgcagtgcttcagccgctaccccgaccacatgaagcagcacgacttcttcaagtccgccatgcccgaaggctacgtccaggagcgcaccatcttcttcaaggacgac
ggcaactacaagacccgcgccgaggtgaagttcgagggcgacaccctggtgaaccgcatcgagctgaagggcatcgacttcaaggaggacggcaacatcctggggcacaagctgg

agtacaactacaacagccacaacgtctatatcatggccgacaagcagaagaacggcatcaaggtgaacttcaagatccgccacaacatcgaggacggcagcgtgcagctcgccgac
cactaccagcagaacacccccatcggcgacggccccgtgctgctgcccgacaaccactacctgagcacccagtccgccctgagcaaagaccccaacgagaagcgcgatcacatggt
cctgctggagttcgtgaccgccgccgggatcactctcggcatggacgagctgtacaagtactcagatctcgagctcaagtagggatcctctagagtcgacctgcagaagcttgcctcgagc
agcgctgctcgagagatctacgggtggcatccctgtgacccctccccagtgcctctcctggccctggaagttgccactccagtgcccaccagccttgtcctaataaaattaagttgcatcatttt
gtctgactaggtgtccttctataatattatggggtggaggggggtggtatggagcaaggggcaagttgggaagacaacctgtagggcctgcggggtctattgggaaccaagctggagtgca
gtggcacaatcttggctcactgcaatctccgcctcctgggttcaagcgattctcctgcctcagcctcccgagttgttgggattccaggcatgcatgaccaggctcagctaatttttgtttttttggtag
agacggggtttcaccatattggccaggctggtctccaactcctaatctcaggtgatctacccaccttggcctcccaaattgctgggattacaggcgtgaaccactgctcccttccctgtccttctg
attttgtaggtaaccacgtgcggaccgagcggccgcaggaacccctagtgatggagttggccactccctctctgcgcgctcgctcgctcactgaggccgggcgaccaaaggtcgcccga
cgcccgggctttgcccgggcggcctcagtgagcgagcgagcgcgcagctgcctgcaggatccgttttgcgctgcttcgcgatgtacgggccagatatacgcgttgacattgattattgacta
ggtcgggcaggaagagggcctatttcccatgattccttcatatttgcatatacgatacaaggctgttagagagataattagaattaatttgactgtaaacacaaagatattagtacaaaatacgt
gacgtagaaagtaataatttcttgggtagtttgcagttttaaaattatgttttaaaatggactatcatatgcttaccgtaacttgaaagtatttcgatttcttggctttatatatcttgtggaaaggacgaa
acaccggaaacctgatcatgtagatcgaacggactctaaatccgttcagccgggttagattcccggggtttccgtttttgctaggtcgggcaggaagagggcctatttcccatgattccttcat
atttgcatatacgatacaaggctgttagagagataattagaattaatttgactgtaaacacaaagatattagtacaaaatacgtgacgtagaaagtaataatttcttgggtagtttgcagttttaa
aattatgttttaaaatggactatcatatgcttaccgtaacttgaaagtatttcgatttcttggctttatatatcttgtggaaaggacgaaacaccggaaacctgatcatgtagatcgaacggactcta
aatccgttcagccgggttagattcccggggtttccgtttttgctaggtcgggcaggaagagggcctatttcccatgattccttcatatttgcatatacgatacaaggctgttagagagataattag
aattaatttgactgtaaacacaaagatattagtacaaaatacgtgacgtagaaagtaataatttcttgggtagtttgcagttttaaaattatgttttaaaatggactatcatatgcttaccgtaacttg
aaagtatttcgatttcttggctttatatatcttgtggaaaggacgaaacaccggaaacctgatcatgtagatcgaacggactctaaatccgttcagccgggttagattcccggggtttccgtttttg
ctaggtcgggcaggaagagggcctatttcccatgattccttcatatttgcatatacgatacaaggctgttagagagataattagaattaatttgactgtaaacacaaagatattagtacaaaata
cgtgacgtagaaagtaataatttcttgggtagtttgcagttttaaaattatgttttaaaatggactatcatatgcttaccgtaacttgaaagtatttcgatttcttggctttatatatcttgtggaaaggac
gaaacaccggaaacctgatcatgtagatcgaacggactctaaatccgttcagccgggttagattcccggggtttccgtttttgctaggtcgggcaggaagagggcctatttcccatgattcctt
catatttgcatatacgatacaaggctgttagagagataattagaattaatttgactgtaaacacaaagatattagtacaaaatacgtgacgtagaaagtaataatttcttgggtagtttgcagtttt
aaaattatgttttaaaatggactatcatatgcttaccgtaacttgaaagtatttcgatttcttggctttatatatcttgtggaaaggacgaaacaccggaaacctgatcatgtagatcgaacggact
ctaaatccgttcagccgggttagattcccggggtttccgtttttgctaggtcgggcaggaagagggcctatttcccatgattccttcatatttgcatatacgatacaaggctgttagagagataatt
agaattaatttgactgtaaacacaaagatattagtacaaaatacgtgacgtagaaagtaataatttcttgggtagtttgcagttttaaaattatgttttaaaatggactatcatatgcttaccgtaact
tgaaagtatttcgatttcttggctttatatatcttgtggaaaggacgaaacaccggaaacctgatcatgtagatcgaacggactctaaatccgttcagccgggttagattcccggggtttccgttttt
tgctaggtcgggcaggaagagggcctatttcccatgattccttcatatttgcatatacgatacaaggctgttagagagataattagaattaatttgactgtaaacacaaagatattagtacaaaa
tacgtgacgtagaaagtaataatttcttgggtagtttgcagttttaaaattatgttttaaaatggactatcatatgcttaccgtaacttgaaagtatttcgatttcttggctttatatatcttgtggaaagga
cgaaacaccggaaacctgatcatgtagatcgaacggactctaaatccgttcagccgggttagattcccggggtttccgtttttgctaggtcgggcaggaagagggcctatttcccatgattcc
ttcatatttgcatatacgatacaaggctgttagagagataattagaattaatttgactgtaaacacaaagatattagtacaaaatacgtgacgtagaaagtaataatttcttgggtagtttgcagttt
taaaattatgttttaaaatggactatcatatgcttaccgtaacttgaaagtatttcgatttcttggctttatatatcttgtggaaaggacgaaacaccggaaacctgatcatgtagatcgaacggact
ctaaatccgttcagccgggttagattcccggggtttccgtttttgctagttattaatagtaatcaattacggggtcattagttcatagcccatatatggagttccgcgttacataacttacggtaaatg
gcccgcctggctgaccgcccaacgacccccgcccattgacgtcaataatgacgtatgttcccatagtaacgccaatagggactttccattgacgtcaatgggtggactatttacggtaaact
gcccacttggcagtacatcaagtgtatcatatgccaagtacgccccctattgacgtcaatgacggtaaatggcccgcctggcattatgcccagtacatgaccttatgggactttcctacttggc
agtacatctacgtattagtcatcgctattaccatggtgatgcggttttggcagtacatcaatgggcgtggatagcggtttgactcacggggatttccaagtctccaccccattgacgtcaatggga
gtttgttttggcaccaaaatcaacgggactttccaaaatgtcgtaacaactccgccccattgacgcaaatgggcggtaggcgtgtacggtgggaggtctatataagcagagctctctggctaa
ctagagaacccactgcttactggcttatcgaaattaatacgactcactatagggagacccaagctggctagcgccaccatggataaaaaaccattagatgttttaatatctgcgaccgggct
ctggatgtccaggactggcacgctccacaaaatcaagcaccatgaggtctcaagaagtaaaatatacattgaaatggcgtgtggagaccatcttgttgtgaataattccaggagttgtagaa
cagccagagcattcagacatcataagtacagaaaaacctgcaaacgatgtagggtttcggacgaggatatcaataattttctcacaagatcaaccgaaagcaaaaacagtgtgaaagtt
agggtagtttctgctccaaaggtcaaaaaagctatgccgaaatcagtttcaagggctccgaagcctctggaaaattctgtttctgcaaaggcatcaacgaacacatccagatctgtaccttcg
cctgcaaaatcaactccaaattcgtctgttcccgcatcggctcctgctccttcacttacaagaagccagcttgatagggttgaggctctcttaagtccagaggataaaatttctctaaatatggca
aagcctttcagggaacttgagcctgaacttgtgacaagaagaaaaaacgattttcagcggctctataccaatgatagagaagactacctcggtaaactcgaacgtgatattacgaaattttc
gtagaccggggttttctggagataaagtctcctatccttattccggcggaatacgtggagagaatgggtattaataatgatactgaactttcaaaacagatcttccgggtggataaaaatctctg
cttgaggccaatgcttgccccgactctttacaactatctgcgaaaactcgataggattttaccaggcccaataaaaattttcgaagtcggaccttgttaccggaaagagtctgacggcaaaga
gcacctggaagaatttactatggtgaacttctgtcagatgggttcgggatgtactcgggaaaatcttgaagctctcatcaaagagtttctggactatctggaaatcgacttcgaaatcgtagga
gattcctgtatggtctttggggatactcttgatataatgcacggggacctggagctttcttcggcagtcgtcgggccagtttctcttgatagagaatggggtattgacaaaccatggataggtgca
ggttttggtcttgaacgcttgctcaaggttatgcacggctttaaaaacattaagagggcatcaaggtccgaatcttactataatgggatttcaaccaatctgtaagaattcaacgcgttaagtcga
ctttaactcgagtctagagggcccgtttaaacccgctgatcagcctcgactgtgccttctagttgccagccatctgttgtttgcccctcccccgtgccttccttgaccctggaaggtgccactccca
ctgtcctttcctaataaaatgaggaaattgcatcgcattgtctgagtaggtgtcattctattctggggggtggggtggggcaggacagcaagggggaggattgggaagacaatagcaggcat
gctggggatgcggtgggctctatggcttctgaggcggaaagaaccctaggggtgcgagcggatcgagcagtgtcgatcactactggaccgcgagctgtgctgcgacccgtgatcttacgg
cattatacgtatgatcggtccacgatcagctagattatctagtcagcttgatgtcatagctgtttcctgaggctcaatactgaccatttaaatcatacctgacctccatagcagaaagtcaaaagc
ctccgaccggaggcttttgacttgatcggcacgtaagaggttccaactttcaccataatgaaataagatcactaccgggcgtattttttgagttatcgagattttcaggagctaaggaagctaaa
atgagccatattcaacgggaaacgtcttgcttgaagccgcgattaaattccaacatggatgctgatttatatgggtataaatgggctcgcgataatgtcgggcaatcaggtgcgacaatctatc
gattgtatgggaagcccgatgcgccagagttgtttctgaaacatggcaaaggtagcgttgccaatgatgttacagatgagatggtcaggctaaactggctgacggaatttatgcctcttccga
ccatcaagcattttatccgtactcctgatgatgcatggttactcaccactgcgatccccagggaaaacagcattccaggtattagaagaatatcctgattcaggtgaaaatattgttgatgcgctg
gcagtgttcctgcgccggttgcattcgattcctgtttgtaattgtccttttaacggcgatcgcgtatttcgtctcgctcaggcgcaatcacgaatgaataacggtttggttggtgcgagtgattttgatg
acgagcgtaatggctggcctgttgaacaagtctggaaagaaatgcataaactcttgccattctcaccggattcagtcgtcactcatggtgatttctcacttgataaccttatttttgacgaggggа
aattaataggttgtattgatgttggacgagtcggaatcgcagaccgataccaggatcttgccatcctatggaactgcctcggtgagttttctccttcattacagaaacggctttttcaaaaatatgg
tattgataatcctgatatgaataaattgcagtttcacttgatgctcgatgagtttttctaatgaggacctaaatgtaatcacctggctcaccttcgggtgggcctttctgcgttgctggcgtttttccata
ggctccgcccccctgacgagcatcacaaaaatcgatgctcaagtcagaggtggcgaaacccgacaggactataaagataccaggcgtttccccctggaagctccctcgtgcgctctcct
gttccgaccctgccgcttaccggatacctgtccgcctttctcccttcgggaagcgtggcgctttctcatagctcacgctgtaggtatctcagttcggtgtaggtcgttcgctccaagctgggctgtgt
gcacgaaccccccgttcagcccgaccgctgcgccttatccggtaactatcgtcttgagtccaacccggtaagacacgacttatcgccactggcagcagccactggtaacaggattagcag

agcgaggtatgtaggcggtgctacagagttcttgaagtggtggcctaactacggctacactagaagaacagtatttggtatctgcgctctgctgaagccagttacctcggaaaaagagttggt
agctcttgatccggcaaacaaaccaccgctggtagcggtggtttttttgtttgcaagcagcagattacgcgcagaaaaaaaggatctcaagaagatcctttgattttctaccgaagaaaggc
cca
